# High-throughput, organ-scale 3D tubule tracking using TubuleMAP

**DOI:** 10.64898/2026.04.23.720231

**Authors:** Chetan Poudel, David Brenes, Wenhui Xie, Yvonne Guan, Pedro Vasquez, Naren Mahadevan, Eesha Konatham, Nina Fan, Ziyu Guo, Nicolas J. Porter, Madeline K. Wong, Michelle M. Martinez, Ruben M. Sandoval, Kenneth W. Dunn, Pierre C. Dagher, Diana G. Eng, Stuart J. Shankland, Jonathan T. C. Liu, Joshua C. Vaughan

**Author notes:** These authors contributed equally to this work (Chetan Poudel, David Brenes, Wenhui Xie).

## Abstract

Advances in tissue clearing and lightsheet microscopy enable mesoscale imaging of intact and convoluted tubular networks, yet analytical tools to map tubule continuity and assess injury patterns within and across tubules are limited. Here, we introduce TubuleMAP, a semi-automated pipeline for 3D tubule tracking and reconstruction that adapts to various morphological and staining patterns, leverages parallel processing of terabyte-scale data for large-scale analysis of tubular networks, and uses a napari interface for human oversight. Using TubuleMAP, we reconstruct 1,000 intact mouse nephrons in ∼1-millimeter-thick kidney slab with ∼400-fold higher throughput and <1% human effort compared to prior approaches. These reconstructions enable analysis of mesoscale nephron organization, quantitative profiling of pathologic morphologies, whole-nephron cytometry, and identification of rare morphologies at unprecedented scales. We demonstrate generalizability by reconstructing all seminiferous tubules in a mouse testis within a day. TubuleMAP is released as an open-source Python package.

## Main

Tubules are fundamental tissue units in many organs, forming continuous epithelial structures that process fluids and transport secretions. In contrast to vascular systems and neuronal networks, epithelial tubules encode physiological function in space: what a cell does depends on its position along the tubule. In the kidney, nephrons consist of a glomerulus that filters the blood followed by a long renal tubule that contains a series of subsegments including the proximal convoluted tubule (PCT), thin limb, thick limb, and distal convoluted tubule (DCT), that progressively modify the filtrate via reabsorption or secretion to maintain fluid and electrolyte homeostasis (1, 2). In the testis, seminiferous tubules form a complex, continuous three-dimensional (3D) network in which germ cells progress through developmental stages, forming a spatial pattern known as the spermatogenic wave, which culminates in the generation of mature sperm (3). Disruption of tubular architecture in both these systems leads to severe disease. In the kidney, renal tubule perturbations have been linked to acute kidney injury, and to chronic kidney diseases such as polycystic kidney disease (4), diabetic nephropathy (5, 6), etc. Injury to seminiferous tubules from impact, torsion, or autoimmune inflammation can result in atrophic or occluded seminiferous tubules, disrupting the spermatogenic wave and negatively impacting fertility (7, 8). Precise morphological characterization and phenotyping of tubular networks in the kidney and testis could provide new insights into disease development and treatment.

Existing technical approaches to study and reconstruct tubular morphology face substantial challenges. Vascular reconstruction methods benefit substantially from vessel sparseness (e.g., vessels occupy only 6% of the renal cortex) and well-established, specific, and convenient labeling strategies such as perfusion or intravenous injection (9). In contrast, epithelial tubules are densely packed (occupying 70% of the renal cortex), are highly convoluted, exhibit ambiguous boundaries, and show segment-specific variation in staining and morphology. Studying an intact functional unit requires following the tubule continuously across large volumes or the entire organ. Individual tubules can be visualized by microdissection or microinjection, but both frequently lose anatomical context, yield incomplete tubules, and are limited in sampling (10, 11). Tubules can be reconstructed through serial histology, a process requiring laborious examination of hundreds of sections and that is prone to tissue distortion and registration errors (12–14). Micro-optical sectioning tomography (15) provides high-resolution datasets with good speed but is destructive to the tissue. Confocal or multiphoton microscopy of cleared tissue (4, 16) produces high-quality, sub-cellular resolution data but remains hindered by slow acquisition speeds. Here, we employed tissue clearing combined with light-sheet microscopy (17) for non-destructive, high-throughput volumetric imaging at sub-cellular resolution. Despite advances in imaging, tubular systems remain difficult to analyze quantitatively. Most studies have been limited to manual centerline tracing (12, 13, 15) and partial 3D segmentation of a few selected segments (4, 18). Large-scale studies of tubule arrangement and morphological alterations with 3D segmentation and reconstructions remain challenging due to the lack of automated, scalable analysis pipelines. While flood-filling networks have worked well for dense 3D reconstructions (e.g., neurons in the brain), they are computationally expensive to run and train (19, 20).

Motivated by these limitations, we developed TubuleMAP, a computationally lean pipeline for semi-automated tubule 3D reconstruction and analysis (e.g., structural metrics and cytometry) that can run on a single GPU-equipped workstation. Instead of performing voxel-wise segmentation across entire organ volumes, TubuleMAP reformulates the problem into a sequence of localized 2D segmentation tasks. It follows each tubule’s centerline and extracts orthogonal cross-sectional planes along its trajectory that can be accurately segmented using general-purpose models such as Cellpose (21, 22). The resulting 2D masks can be interpolated to reconstruct the full tubule in 3D. To improve robustness to morphological and staining variations, a model-switching module automatically selects the most suitable segmentation model on-the-fly as cross-sections are processed. An interactive human-in-the-loop interface enables guided exploration and rapid correction when rare segmentation errors occur. TubuleMAP is designed to handle terabyte-scale datasets via chunked data formats and scalable parallel processing. Through validation on thick mouse kidney slabs and whole mouse testis, we demonstrate that TubuleMAP provides unprecedented scale in 3D reconstruction of tubules, enabling high-throughput and statistically robust investigations of tubule pathologies (hundreds to thousands of units) at ∼1% of the human effort required by manual approaches.

## Results

### TubuleMAP pipeline

In the kidney, analysis of whole nephrons requires uniform labeling across large tissue volumes, given that nephrons extend millimeters and span multiple regions, from the outer cortex through the juxtamedulla into the deep medulla, and transition through distinct segments, including the PCT, thin limb, thick limb, and DCT. The thin limbs can taper to ∼15 µm in diameter and are densely packed in the inner medulla, placing stringent demands on imaging resolution. Our workflow uses small-molecule, 3-channel FLARE (Fluorescence Labeling of Abundant Reactive Entities) staining to rapidly and uniformly label carbohydrates, protein amines, and DNA throughout millimeter-thick tissue slabs with a high signal-to-noise ratio (23, 24). Tissues were cleared using organic solvents and ethyl cinnamate (25), then imaged by OTLS microscopy (17) at high resolution (**Supplementary Video 1**). After image post-processing, the volume is stored as a multi-resolution Zarr pyramid (OME-Zarr (26)) for random access to small data chunks, which makes image processing scalable. TubuleMAP utilizes the carbohydrate channel to perform parallelized 3D nephron reconstruction with infrequent human-in-the-loop assistance, enabling comprehensive downstream analysis (**Fig. 1a**). TubuleMAP is implemented as an open-source Python software package for 3D tubule tracking and segmentation that is now available on GitHub (see Code Availability).

**Figure 1.**
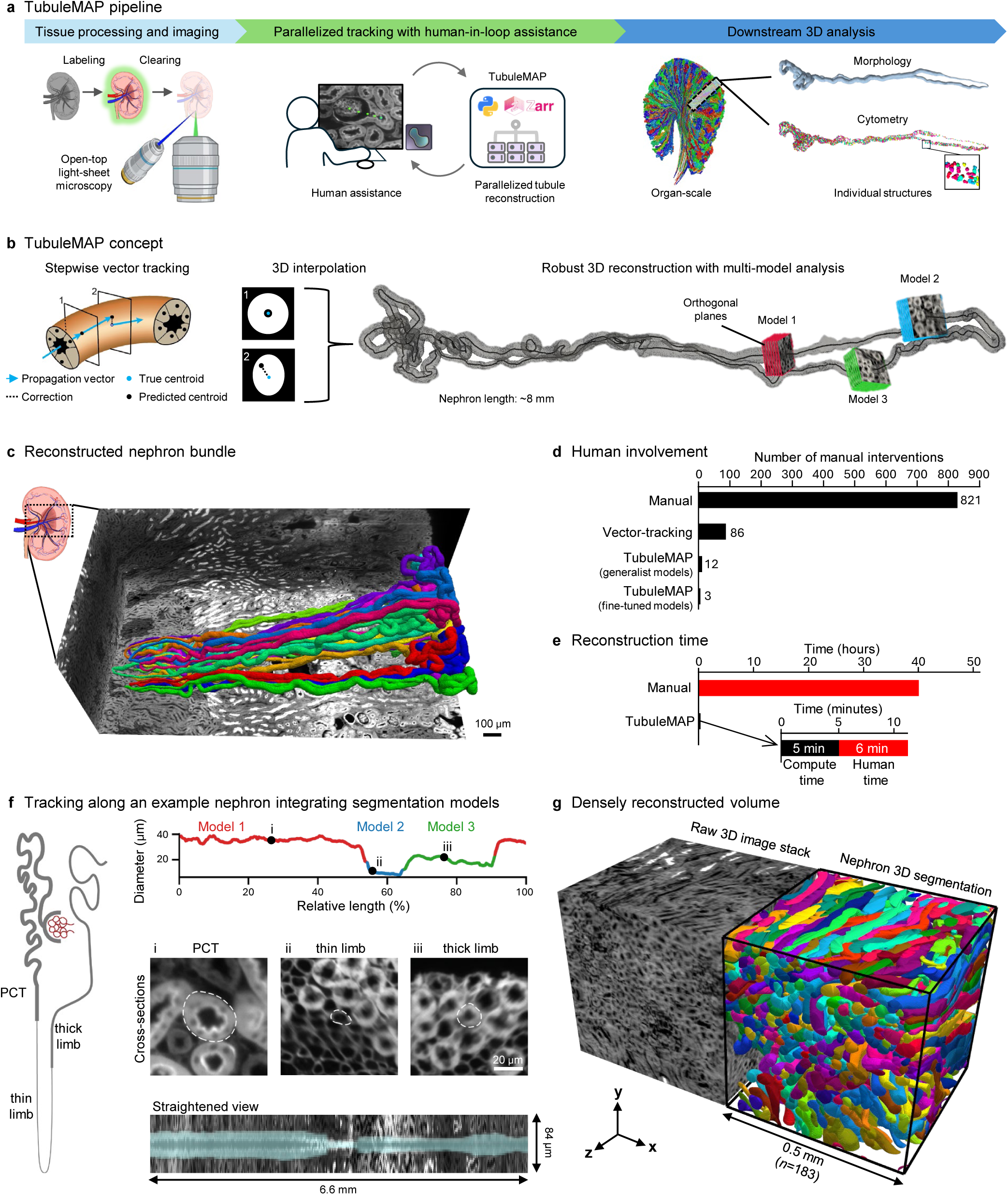
TubuleMAP overview and performance. (**a**) End-to-end pipeline from tissue processing and imaging to high-throughput tubule reconstruction and downstream quantitative analysis. (**b**) TubuleMAP propagates the centerline by stepwise vector tracking, correcting its trajectory using segmentations of orthogonal cross**-**sections. These cross-sections are interpolated to generate a 3D reconstruction of the tubule. A model-switching module automatically selects the best-performing segmentation model for different nephron segments. (**c**) FLARE-stained kidney volume with 20 manually traced nephrons used as ground truth for benchmarking. (**d**) Mean number of human interventions per nephron across reconstruction methods. (**e**) Reconstruction time per nephron for manual versus TubuleMAP. (**f**) Example segment-specific model switching; color transitions mark model-switch events along the nephron. Representative orthogonal cross-sections (i–iii) and a morphologically straightened view with overlayed segmentation are shown below. (**g**) Reconstruction in a dense kidney region for a cube of 0.5 mm edge length.

Starting from two user-defined seed points, TubuleMAP performs stepwise vector tracking to follow the tubule centerline (**Fig. 1b**). At each step, the current propagation direction is used to extrapolate the next centerline position, defining a predicted centroid. A plane orthogonal to the propagation direction and centered on the predicted centroid is sampled and segmented. TubuleMAP computes the true centroid from the segmentation mask; the offset between the predicted and true centroids is used to update the propagation vector. Repeating this procedure yields the tubule centerline and a sequence of cross-sections along the tubule, which are then interpolated to form a continuous 3D tubule reconstruction (**Supplementary Video 2**).

Segmentation models sometimes fail to generate a cross-section mask due to variations in tissue physiology, staining, imaging conditions, or spatial orientation (**Supplementary Fig. S1)**. For example, generalist segmentation models such as Cellpose reliably produce masks for round or elliptical cell-like shapes but often fail to segment elongated tube-like structures. To address this failure and avoid interruptions of the tracking process, we created three TubuleMAP modules (described in greater detail in **Supplementary Fig. S2**) that are invoked to overcome mask failures: *(i) Rotational search* tilts the sampling plane across the tubule over a discrete set of angles to identify the view and propagation direction that minimizes eccentricity. This helps overcome sharp turns in convoluted tubules. *(ii) Adaptive parameter scaling* calculates the tubule diameter from recent cross-sections and uses it to continuously tune size-specific priors for tracking and segmentation. This allows for adapting to gradual changes in tubule diameter, when tubules narrow or widen at organ-level transitions or in disease. *(iii) Troubleshooting and model switching* modules overcome abrupt changes in morphology or staining patterns by leveraging 3D multi-hypotheses, where masks generated from several segmentation models and diameter combinations are integrated into one consensus segmentation using Ultrack (27). Each hypothesis is then compared to the consensus; the highest-agreement hypothesis (i.e., model and associated diameter) is adopted for the next iteration.

In rare cases where a mask is not generated by any of the models, tracks are automatically flagged and queued into an interactive, human-in-the-loop interface for manual intervention. This enables a user to serially correct tracks with minimal effort using one or a few point clicks, then reassign them to the tracking queue to resume automated, parallelized reconstruction (**Supplementary Video 3**). Further details on the user interface are provided in **Supplementary Fig. S3** and **Supplementary Video 4.**

### Performance assessment in the mouse kidney

Tracking performance was benchmarked in 20 manually annotated nephron tracks (∼150 mm total tubule length) from previously published data (24) (**Fig. 1c**). Compared to manual tracking, stepwise vector tracking with a generalist model (Cellpose ‘cyto2’) reduced the average number of human interventions per nephron from 821 to 86. Incorporating all three TubuleMAP modules further reduced the number of interventions to 12. Ablation studies (**Supplementary Fig. S4a(i)**) confirmed that the troubleshooting module produced the largest reduction in human interventions across all segments (PCT, thin limb, and thick limb), highlighting that recovery via multi-hypothesis integration across different diameters is crucial. Adaptive parameter scaling yielded the next-largest improvement, while rotational search provided smaller, additional segment-dependent improvements. When TubuleMAP had access to a suite of three fine-tuned, segment-specific models, it showed the best performance, with only 3 human interventions (∼5 minutes of human time) required per nephron (**Fig. 1d-f; Supplementary Fig. S4a(ii), S4b(ii)**). While rotational search and troubleshooting via model switching do improve performance, they are more computationally expensive than the standard stepwise tracking (∼10× and ∼100× slower, respectively; **Supplementary Fig. S4c**). To maintain code efficiency, TubuleMAP triggers rotational search only for highly elongated cross-sections and triggers troubleshooting only when the current model fails to generate a mask. Resolution also plays a key role in tracking performance. Using 2× and 3× down-sampled volumes derived from the original dataset, tracking performance degraded sharply (**Supplementary Fig. S4d**). The nephron’s thin limb had the shortest uninterrupted length tracked, reflecting that it was most difficult to reconstruct due to its small diameter (∼15 µm) and densely packed arrangement.

TubuleMAP’s 3D segmentation performance was evaluated by comparing against two manually segmented nephrons. The intersection over union was 0.73 and 0.75 (**Supplementary Fig. S5**). Manual 3D segmentation required ∼40 human hours per nephron, whereas TubuleMAP achieved comparable reconstructions in ∼11 minutes per nephron through parallelization (5 minutes of compute time and 6 minutes of human time; **Fig. 1d,e** and **Supplementary Fig. S4e**). This corresponds to <1% of the human effort and an approximately 400-fold increase in throughput compared to manual approaches. **Fig. 1f** shows an example of a ground truth nephron reconstructed (tracked and segmented) by TubuleMAP and illustrates how TubuleMAP adjusts its tracking parameters (diameter) and switches between segment-specific models across the PCT, thin limb, and thick limb. The straightened view projects the nephron along the centerline, making transitions between segments and other morphological features easy to visualize. **Fig. 1g** highlights the high packing density of renal tubules in the mouse kidney: within a 0.5 mm × 0.5 mm × 0.5 mm cube, 183 different nephrons were tracked. Despite this high density, TubuleMAP performs robust reconstructions.

#### Case study: TubuleMAP analysis of 1000 nephrons in a single mouse kidney tissue slab

We applied TubuleMAP to reconstruct 1000 nephrons in 3D within a 1.3 mm-thick slab of healthy mouse kidney tissue (**Fig. 2a, Supplementary Video 5**), representing an over 100-fold increase in scale compared to previous studies with only partial, manual 3D segmentations (4). Each nephron was tracked from the glomerulus through the proximal convoluted tubule, thin limb, thick limb, and back to the glomerulus contact point at the macula densa. This structural feature of nephrons, whereby the tubule contacts the glomerulus from which it originated after many millimeters of travel, provides an internal anatomical validation of tracking accuracy. To enable region-aware analysis of nephron morphology, we trained an nnU-Net model to divide the volume into four regions with high accuracy based on kidney-specific staining patterns: cortex, outer stripe of outer medulla (OSOM), inner stripe of outer medulla (ISOM), and inner medulla (IM) (**Fig. 2b, Supplementary Fig. S6**). Inspired by the Allen Mouse Brain common coordinate framework (28), this 3D regional atlas provides organ-scale spatial context for each reconstruction.

**Figure 2.**
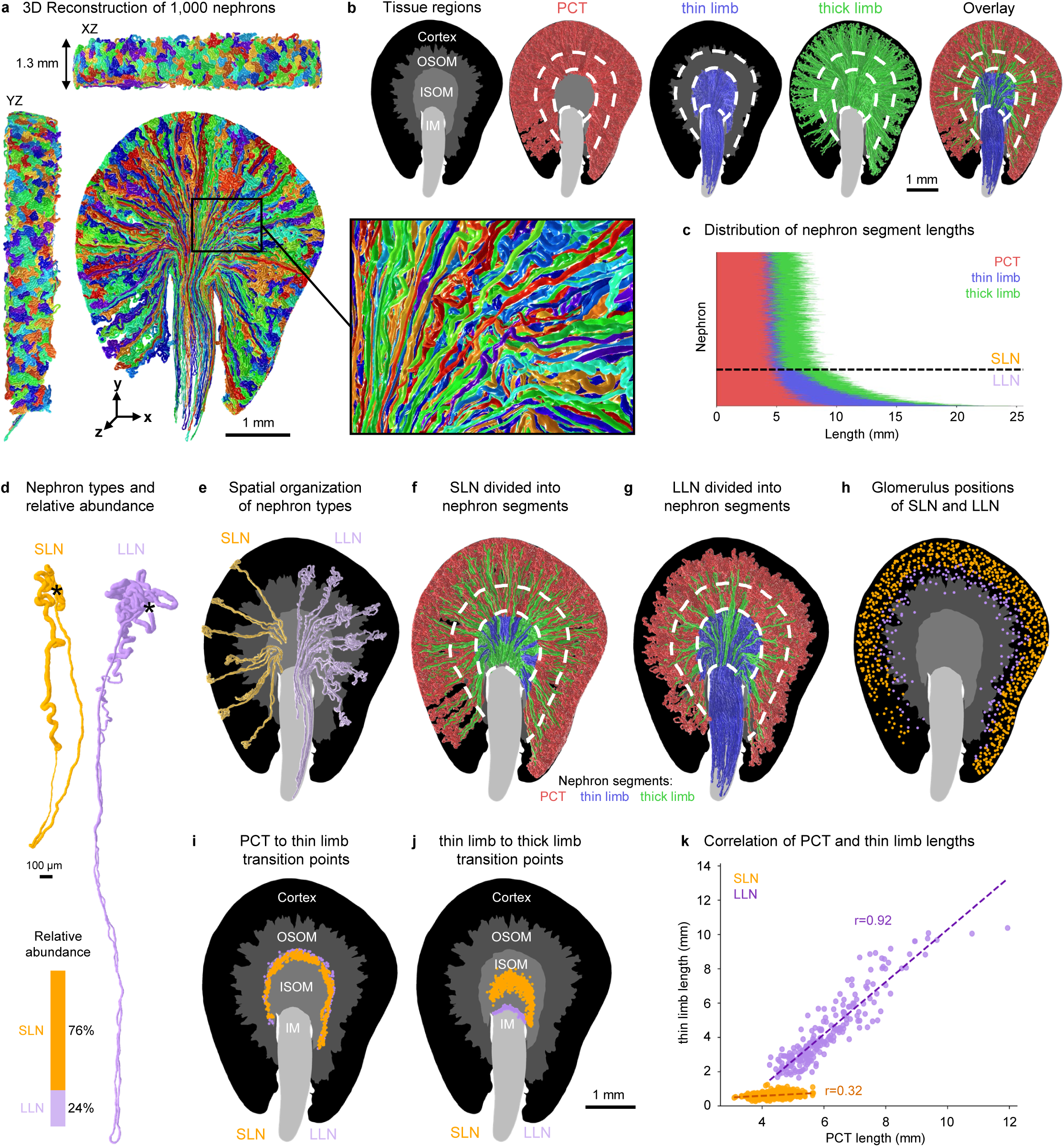
Mesoscale organization of nephron architecture in the mouse kidney. (**a**) 3D reconstruction of 1000 nephrons in a 1.3 mm thick mouse kidney slab. (**b**) Nephron segments (proximal tubule (PCT), thin limb, thick limb) are shown in an organ-scale context using a four-region atlas: cortex, outer stripe of the outer medulla (OSOM), inner stripe of the inner medulla (ISOM), and inner medulla (IM). (**c**) Distribution of nephron segment lengths, highlighting thin limb length as a basis to distinguish short-loop (SLN) from long-loop (LLN) nephrons. (**d**) Visual differences and relative abundance of SLNs and LLNs, with glomerulus positions shown using *. (**e-h**) Differences in the spatial organization of SLNs and LLNs, shown using segment location and glomerulus location with respect to tissue regions. (**i-j**) Locations of segment transitions by nephron type. (**k**) PCT segment grows proportionally with thin limb length in LLNs but not in SLNs.

Reconstructed nephrons were manually subdivided into the PCT, thin limb, and thick limb segments based on nephron morphology. The transitions between segments were abrupt, generally corresponding to transitions identified in the 3D regional atlas (**Fig. 2b**), with distinct average diameters for each segment (PCT = 43 ± 2 µm, thin limb = 16 ± 5 µm, thick limb = 24 ± 3 µm, mean ± SD; **Supplementary Fig. S7a**). PCTs were the longest segment (**Fig. 2c**) and displayed the greatest curvature at the outer cortex (**Supplementary Fig. S7b-c**). Curvature and torsion were both high in the outer medulla region due to sharp turns at the loop of Henle. Consistent with past literature discussing nephron types (2, 29, 30), we identified two distinct populations of nephrons that correspond to short-loop nephrons (SLNs, ∼76% abundance) and long-loop nephrons (LLNs, ∼24% abundance) (**Fig. 2d**). We found that SLNs and LLNs could be cleanly distinguished based on thin limb length (<1.4mm for SLN, **Fig. 2c, Supplementary Fig. S7d**) or, equivalently, by whether the nephron entered the inner medulla region (**Fig. 2e-g, Supplementary Video 6**). In contrast, the distance from a nephron’s glomerulus to the renal capsule, which is a commonly used classification metric, was only moderately predictive of nephron type, achieving at best 90% accuracy at a depth threshold of 330 µm (**Fig. 2h**).

Segment transitions occurred at consistent positions along the corticomedullary axis. As expected, the PCT-to-thin limb transitions clustered near the OSOM-to-ISOM boundary for both SLN and LLN (**Fig. 2i**). The thin-to-thick limb transitions were in the middle of the ISOM for SLN and in the ISOM-to-IM boundary for LLN (**Fig. 2j**). Among the SLNs, roughly half (390/762) lacked an ascending thin limb, consisting of only a descending thin limb that transitions into a thick limb before the loop of Henle (**Supplementary Fig. S8**). The remaining SLNs had both descending and ascending thin limbs, similar to LLNs. Segment lengths also scaled differently across nephron types, where thin limb length is strongly correlated with PCT length for LLNs (r = 0.9) but only weakly correlated in SLNs (r = 0.32). **(Fig. 2k**) This recapitulates the developmental differences between the nephron types: the early-developing LLNs and late-developing SLNs. The segments of LLNs likely grow proportionally during early kidney development towards the medulla, while the growth of thin limbs in SLNs is restricted in late kidney development due to crowding. Additional correlations between segment lengths are shown in **Supplementary Fig. S7e**. After sampling 1000 nephrons, ∼1% of nephrons with atypical morphologies were encountered and manually confirmed. These nephrons included no thin limbs or were confined entirely to the cortex (**Supplementary Fig. S9)**.

#### Individual nephron morphology analysis in pathology

In addition to quantifying the mesoscale spatial arrangements of tubule networks, TubuleMAP includes a novel suite of computational tools for downstream analysis. For each reconstruction, spatially resolved morphometric profiles (e.g., diameter, curvature, and torsion) were computed along the full tubule. These measurements enable both visualization and quantification of tubular remodeling. For example, in **Fig. 3a**, a representative atrophied renal tubule in a mouse model of focal segmental glomerulosclerosis (FSGS) disease shows reduced diameter and higher local curvature, while a dilated or hypertrophied tubule in an aged mouse shows larger diameter and lower curvature relative to a healthy nephron. The reconstructions also enable mapping of glomerulo-tubular injury patterns through direct comparison of glomerular appearance and tubule morphology (**Fig. 3b**). TubuleMAP also includes a morphological straightening procedure, which transforms complex 3D tubular structures into simplified 2D visualizations (longitudinal cut-through views), providing an intuitive snapshot of the nephron morphology and facilitating rapid visual assessments and comparisons across nephrons (**Fig. 3c**). Using the manually assigned nephron segment labels, the same morphometric profiles can also be summarized by subsegment (PCT, thin limb, thick limb), allowing direct comparisons of segment-specific features across experimental conditions and disease states.

**Figure 3.**
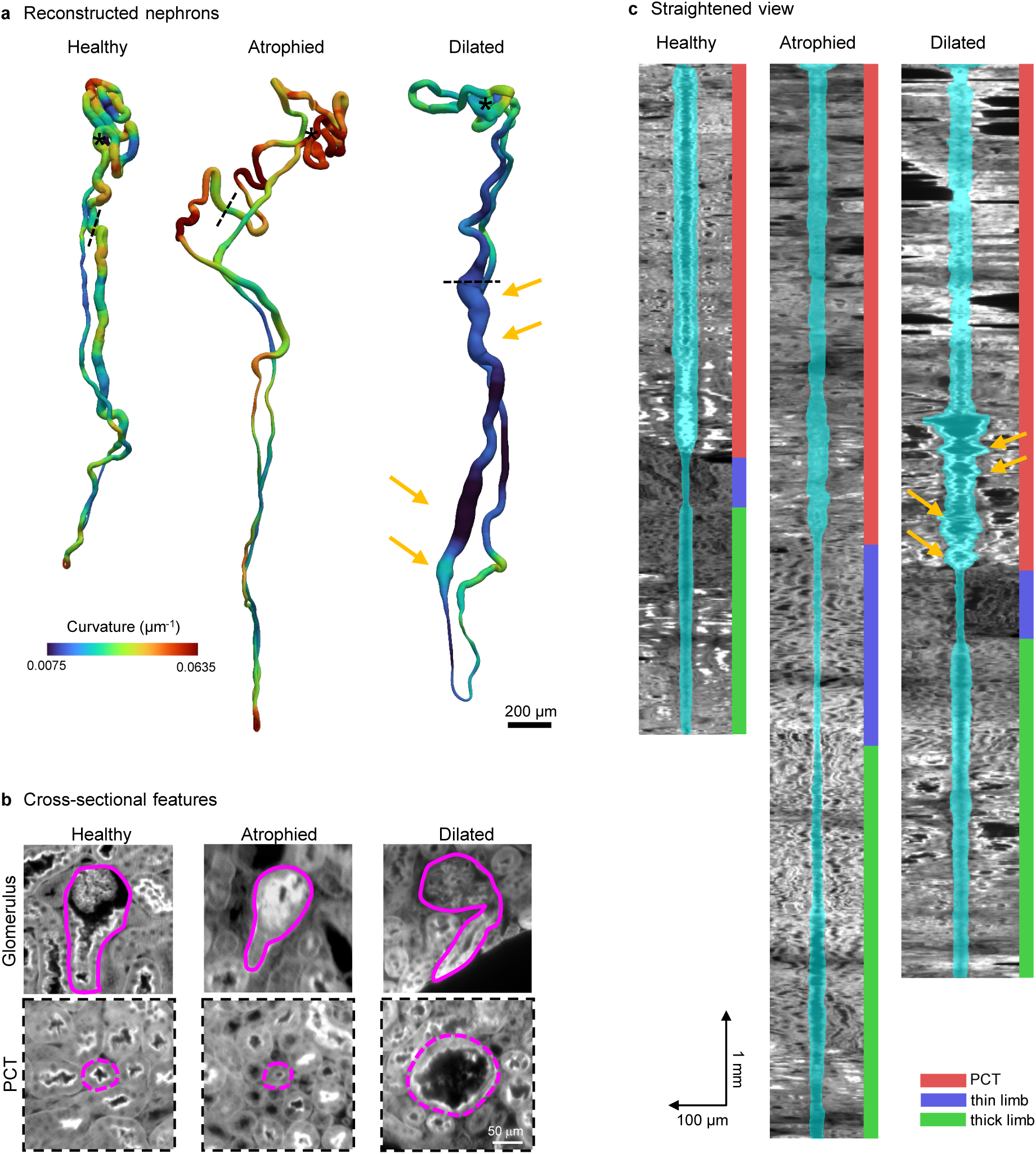
Single nephron analysis of pathologic morphologies. (**a**) Representative reconstructed nephrons showing normal, atrophied, and dilated tubules, color-coded by local curvature. (**b**) Cross-sectional views of the glomerulus and the proximal tubule (PCT) of the reconstructed nephrons in (a); the locations of the PCT planes are indicated by dashed lines in (a). (**c**) Straightened views of nephrons in (a) with the segmentation shown in cyan and segment identities indicated by color bars.

#### Nephron cytometry

Using the 3D reconstructions, the DNA channel was masked to extract nuclear information for individual nephrons. To our knowledge, these are the first cytometry datasets collected and analyzed at the whole nephron scale (**Supplementary Video 7**). While DNA staining was not uniform throughout the 1.3 mm thickness, we focused on 20 nephrons with complete labeling. **Fig. 4a** shows representative nephron nuclei segmentations for a SLN and a LLN, while **Fig. 4b** provides higher-magnification views highlighting nuclei organization in the PCT, thin limb, and thick limb. LLNs occupy more volume and have more nuclei (3803 ± 1243, mean ± S.D., n=10) than SLNs (2926 ± 260, mean ± S.D., n=10). In both nephron types, the PCT segment accounted for 75-80% of the total nuclei count. Although segment diameters (**Fig. 4c**) and nuclei densities along the length (**Fig. 4d**) differed substantially across nephron segments, these differences were largely erased when volumetric nuclei densities were computed (**Fig. 4f**).

**Figure 4.**
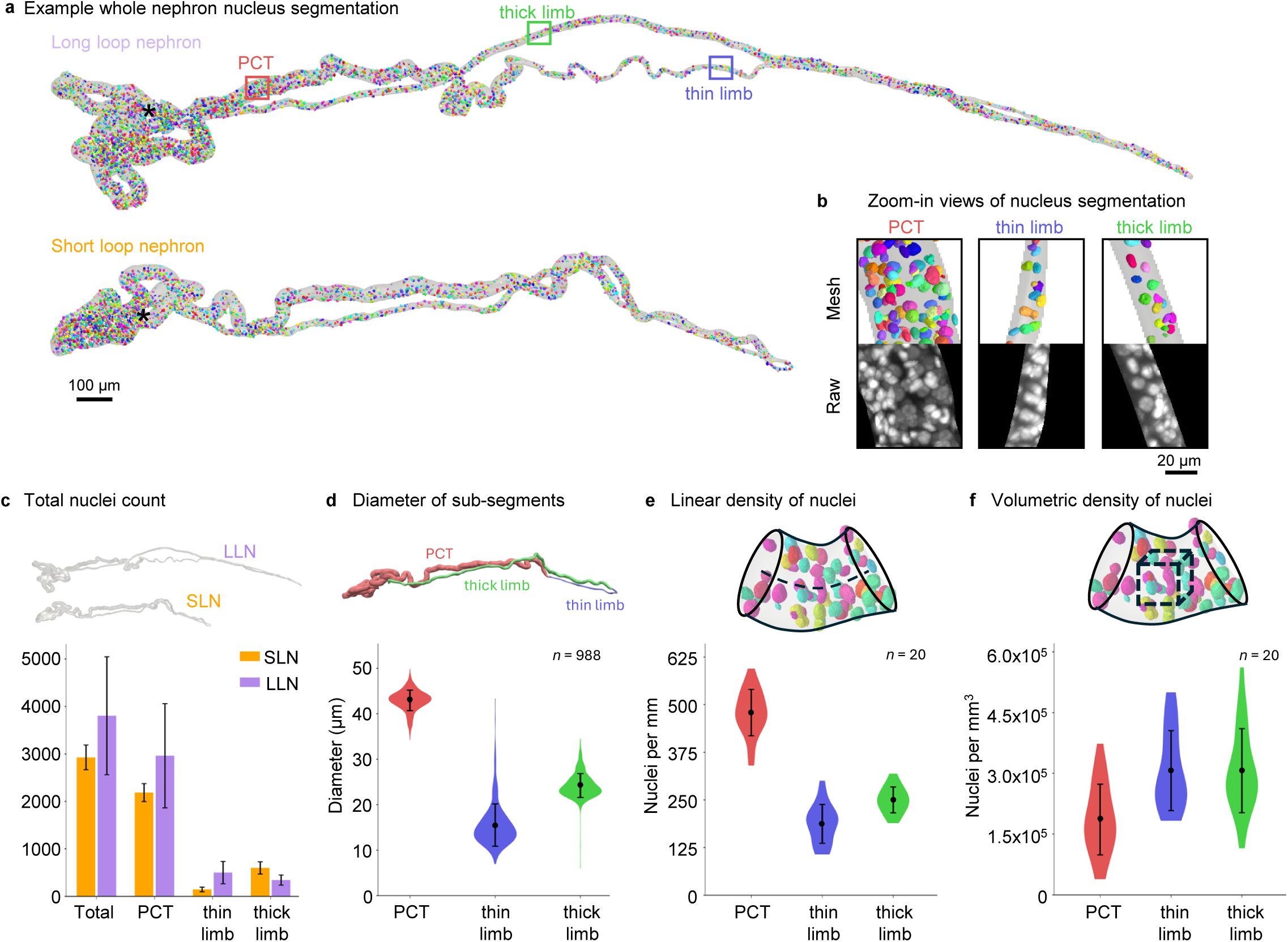
Whole–nephron cytometry. (**a**) Example long loop and short loop nephrons showing reconstruction outline and segmented nuclei. (**b**) High-magnification views of regions highlighted in (a) showing the raw nuclei channel data and nuclei segmentations for each segment. (**c**) Total nuclei counts for LLN and SLN and sub-divided by segments. For all nephrons, (**d**) diameter of segments, (**e**) linear nuclei density, and (**f**) volumetric nuclei density are plotted. Mean +/- S.D. is shown

#### Generalizability to other tubular structures

We demonstrate the generalizability of TubuleMAP to segment other tubular structures by mapping all the seminiferous tubules in a mouse testis. The intact mouse testis (10.1 mm × 5.4 mm × 4.6 mm) was labeled via FLARE, cleared using CUBIC protocols (31), and imaged using the OTLS microscope. TubuleMAP with a generalist model (Cellpose ‘cyto3’) was applied to the carbohydrate channel to reconstruct all 13 seminiferous tubules, totaling ∼2.6 m of length, within 1 day. This process was largely automated, requiring only a few hours (∼5 hours) of manual oversight, and could be further improved by developing testis-specific segmentation models. **Fig 5a** and **Supplementary Video 8** show 3D renderings of all seminiferous tubules in different colors, and **Fig. 5b** shows each individual tubule in the same reference view, highlighting their highly coiled organization and distinct spatial distributions within the testis.

**Figure 5:**
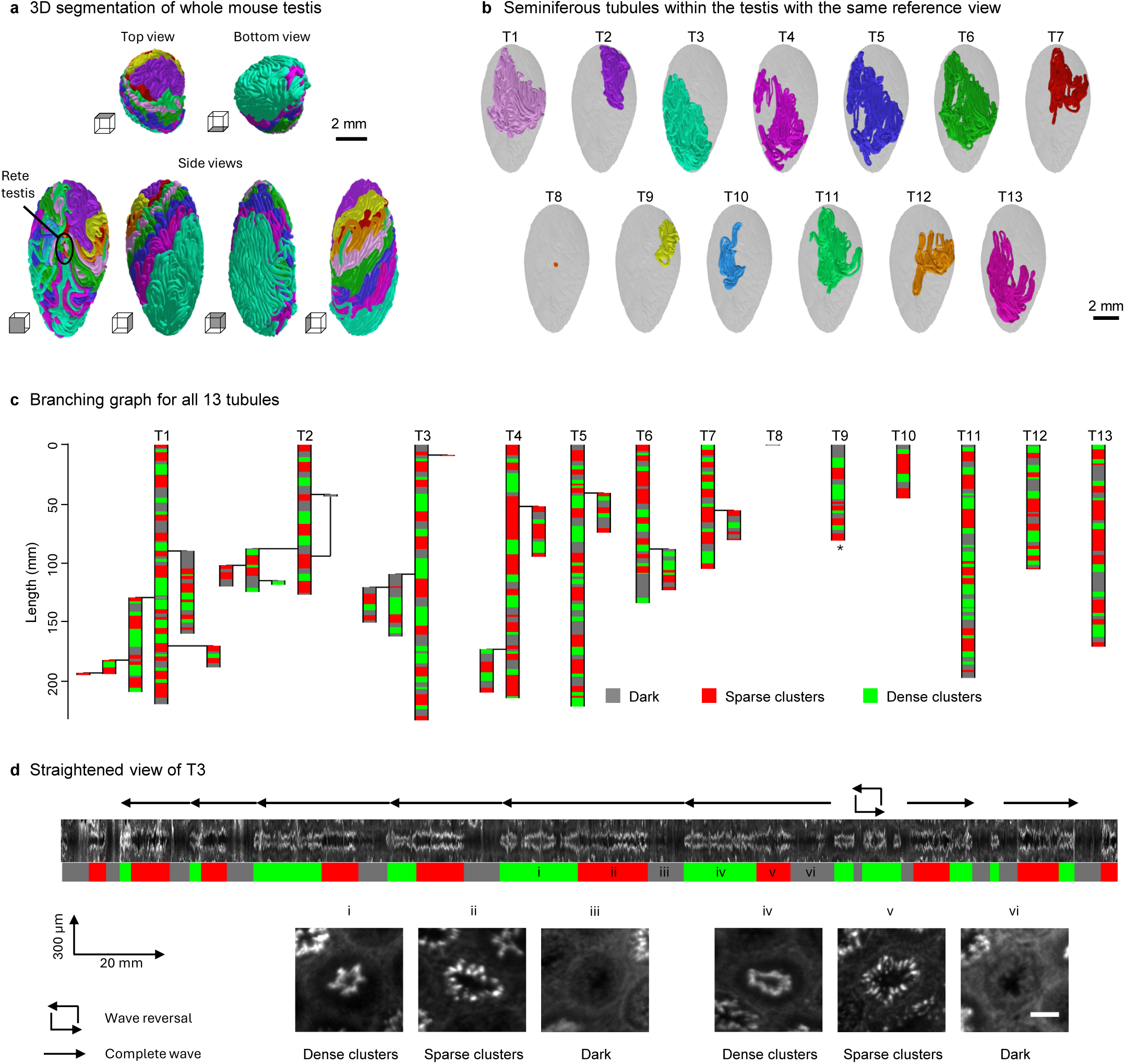
Applying TubuleMAP to fully segment and study the whole mouse testis. (**a**) 3D reconstructions of a whole mouse testis, shown at different orientations and with each of the 13 seminiferous tubules assigned a different color. (**b**) Spatial orientation of the 13 individual seminiferous tubules seen from the same perspective. (**c**) Graph showing tubule branching structure and classification by spermatogenic wave pattern. The longest branch is shown as the main tubule. Unless marked by an asterisk (*), all tubule ends are located in the rete testis. (**d**) Straightened view of T3, with periodic nuclear signal patterns in the DNA channel serving as a proxy for identifying spermatogenic waves. The colored bars below the straightened view shows the three classified spermatogenic stages (green, dense clusters class; red, sparse clusters class; blue, dark class). Tubule cross-sections i-vi show DNA staining patterns at different stages of the spermatogenic wave. A wave reversal is noted in the tubule in which the class sequence changed direction. Scale bar: 75 µm.

Unlike nephrons, seminiferous tubules frequently exhibit branching; branch points were automatically identified during the reconstruction (see Methods). **Fig. 5c** summarizes the branching architecture of all 13 seminiferous tubules. 7 tubules contained 17 total branches. There were 12 primary branches originating from the main tubule, 4 secondary branches, and 1 tertiary branch. In one case (T2), a branch later merged with the main tubule; this reconnection event was not counted as an additional branching point. All but one tubule ended at the rete testis; one end of T9 was a blind end, meaning that the tubule abruptly stopped. The 2-month-old mouse likely had not reached sexual maturity, as one of the seminiferous tubules (T8) was still underdeveloped and short.

For each seminiferous tubule, morphological straightening was applied to examine the variation in the DNA staining pattern along its length (**Fig. 5d**). Visual inspection revealed three distinct patterns (dark, sparse clusters, and dense clusters) that together form a period of the spermatogenic wave (**Fig. 5d**); high-resolution examples of these classes are shown in **Supplementary Fig. S10a and Supplementary Video 9**. These three classes roughly align with the twelve stages (I–XII) of the spermatogenic wave identified in previous studies (32). The dense clusters class corresponds to stages I–V and X–XII, the sparse clusters class corresponds to stages VI–VII, and the dark class corresponds to stages VIII–IX. We then developed an automated algorithm to classify cross-section images into these classes (see Methods; **Supplementary Fig. S10b**) and applied it to all tubules. Overall, 71 complete spermatogenic waves were detected. These waves measured 18.2 ± 6.3 mm in length on average, matching a prior study in young adult mice reporting 76 complete waves with an average length of 16.8 mm (33). In **Fig. 5c**, tubules are color-coded according to this classification. Across all tubules, 29%, 40%, and 31% of cross-sections were dark, sparse clusters, and dense clusters, respectively. **Fig. 5d** shows the straightened view of the longest, continuous tubule in T3 and highlights 8 distinct spermatogenic periods, including one reversal in which the class sequence changed direction.

## Discussion

Rapid advancements in mesoscale labeling, tissue clearing, and volumetric imaging have enabled the capture of large organ-scale datasets with subcellular detail. However, the quantitative analysis of dense, long, and highly convoluted tubular structures (e.g. nephrons, collecting ducts, seminiferous tubules, etc.) has until now remained a major barrier to understanding organ-scale architecture and changes in disease progression. TubuleMAP overcomes this barrier by replacing intractable dense 3D segmentation with a fundamentally more scalable tracking-and-reconstruction strategy. TubuleMAP follows each tubule through sequential 2D cross-sectional segmentations and integrates these into complete 3D reconstructions. Because tubules are reconstructed independently, the workflow is well-suited for parallelization, enabling efficient and high-throughput analysis of large tubule populations within intact organs.

By integrating multiple modules to improve robustness and reduce the need for human intervention, TubuleMAP enables research studies at a scale that was previously impractical or unattainable. For example, a modest experimental design using three biological replicates per condition (health and disease) with 100 nephrons per sample, would require an estimated 11 human years of manual segmentation but can be completed with ∼60 hours of human effort using TubuleMAP. Notably, TubuleMAP reconstruction of nephron proximal tubules often proceeds with no human correction. Since the PCT is most prone to acute and chronic injury (34), the ability to perform population-level analyses of this nephron segment will advance the field. Furthermore, whole-nephron reconstructions create new opportunities to study segment-specific growth, injury, and remodeling during development, disease, and aging. Critically, they also provide access to complex 3D relationships between proximal and distal segments of a nephron that underlie feedback mechanisms regulating glomerular function. These relationships are invisible in conventional 2D analyses and are frequently disrupted in disease. Beyond individual nephrons, TubuleMAP enables mapping of spatially coordinated changes and remodeling across neighboring tubules, facilitating studies of how local perturbations propagate through tissue networks. For example, TubuleMAP could be used to map the effects of collecting duct obstruction on connected nephrons in kidney stone disease or to relate local vascular injury to adjacent tubular damage in diabetic kidney disease. Additionally, TubuleMAP enables cytometry along the nephron axis which could be applied to study kidney diseases where tubular injuries lead to cellular loss in segment-specific patterns.

The same framework generalizes naturally to other tubular systems. Studying the 3D morphological changes within and along seminiferous tubules in the testis could further our understanding of male reproductive health and disease. TubuleMAP enables reconstruction of seminiferous tubules directly from volumetric data, overcoming the limitations of prior approaches based on sparse, serial histological sections that were primarily used to characterize global tubular architecture (33, 35–37). This allows consistent, accurate, and near-isotropic quantification of tubule features such as diameter, curvature, and torsion, as well as cellular organization along the tubule axis. The ability to generate straightened views of entire tubules provides an especially powerful new mode of visualization, enabling rapid identification of subtle histological variations such as spermatogenic wave states. While the testis datasets analyzed here were acquired at relatively modest resolution, the approach is immediately compatible with higher-resolution datasets like ones used in our kidney study. Applied at such scales, TubuleMAP has the potential to resolve the full spectrum of 12 developmental stages of mouse spermatogenesis described in prior studies (33).

TubuleMAP’s performance depends on accurate segmentation at each step. Most segmentation errors that interrupt tracking arose from local imaging defects such as tile striping artifacts, blur from incomplete tissue clearing, or sample damage. Moving the workflow toward near-full automation would require improving image quality and improving the algorithms used to segment each cross-section. While TubuleMAP is readily compatible with a wide range of organ-scale imaging modalities including confocal microscopy, lightsheet microscopy, micro-optical sectioning tomography (15), x-ray phase-contrast tomography (38), etc., the evolution of these technologies will lead to enhanced image quality and corresponding gains in segmentation and automation. Image resolution can also be improved with expansion microscopy (39–41). On the algorithmic side, segmentation accuracy can be improved using post-processing techniques like denoising and deconvolution (integrated into Cellpose v3 (42)), or by incorporating additional deep-learning architectures and models (e.g., combining Cellpose and Segment Anything for Microscopy (43)). Even with these advances, occasional segmentation errors will persist in the most challenging regions. As in other large-scale reconstruction efforts, the practical goal is to confine human intervention to these rare edge cases rather than to eliminate them entirely. Human verification will remain valuable for quality assurance. TubuleMAP is explicitly designed around this principle, with an interactive interface for rapid human review and correction where needed, ensuring both scalability and reliability.

The required level of reconstruction fidelity will depend on the biological application. Many architectural and morphometric analyses remain robust to minor local errors, provided that major shape trends and tubule trajectory are preserved. In contrast, applications such as nephron cytometry demand higher geometric precision, particularly when assigning nuclei or other cellular features to specific nephrons in densely packed regions. Because no prior method has achieved whole nephron reconstructions at this scale, it remains difficult to define a universal threshold or benchmark for sufficient reconstruction accuracy. Nevertheless, the accuracy achieved here demonstrates that TubuleMAP reaches a level sufficient for a wide range of quantitative analyses, while establishing a foundation for further refinement as imaging and segmentation technologies evolve.

By enabling comprehensive 3D reconstructions at organ scale, TubuleMAP opens an entirely new analytical space for tubule studies. These data make it possible to quantify spatial injury patterns within individual tubules, across neighboring structures, or along connected networks. Future work could use these reconstructions to study inter-tubule crosstalk, perform computational modeling of fluid flow and transport kinetics, and guide the fabrication of bioprinted tissue models (44).

Taken together, TubuleMAP establishes a powerful and extensible framework for mapping tubular architecture with unprecedented scale and resolution. By converting a longstanding technical barrier into a tractable computational problem, it enables a new class of studies that bridge cellular, structural, and organ-level biology. This capability positions TubuleMAP to fundamentally reshape how tubular systems are analyzed, accelerating discovery across physiology, development, and disease.

## Online Methods

### Chemicals and reagents

Fluorescent dyes were purchased as follows: SYBR Green I (Invitrogen, S7563), ATTO 565 hydrazide (ATTO-TEC GmbH, AD 565-121), ATTO 647N NHS ester (Sigma-Aldrich, 18373-1MG-F). General reagents were purchased as follows: Phosphate-Buffered Saline (PBS; Invitrogen, 10010049), sodium azide (NaN_3_; S2002), 32% paraformaldehyde (PFA) aqueous solution (Electron Microscopy Sciences, RT15714). FLARE staining used sodium periodate (NaIO_4_; Millipore Sigma 311448), sodium cyanoborohydride (NaCNBH_3_; 156159), sodium acetate buffer (NaOAc; Fisher Scientific, cat. no. S209), sodium cyanoborohydride (NaCNBH_4_; Millipore Sigma, 156159-10G) and 2-(N-Morpholino) ethanesulfonic acid (MES buffer; AK Scientific Product Catalog, M3671-50G). The tissue clearing process used 100% Ethanol (Fisher Scientific, 04-355-450) or tetrahydrofuran (THF; Thermo Fisher, 030760.K2), dichloromethane (Sigma-Aldrich, 75-09-2), and ethyl cinnamate (ECi; MilliporeSigma, W243000-1KG-K).

### Tissue preparation: labeling and clearing

All protocols and methods involving animals in this work were approved by the Institutional Animal Care and Use Committee at Indiana University School of Medicine and University of Washington. Four male C57BL/6 mice were used in this study, including three untreated mice (ages 2, 4, and 27 months). The fourth mouse, aged 3.5 months (**Fig. 3**, “atrophy”), was administered 2 consecutive doses of anti-podocyte antibody at 10 mg / 20 g bodyweight and euthanized after 14 days to induce an experimental model of focal segmental glomerulosclerosis (FSGS). Mice were anesthetized by isoflurane/oxygen mixture followed by cardiovascular perfusion with 1× PBS for 5 min and then 4% PFA solution in 1× PBS for 5 min. Kidneys or testes were collected and drop-fixed in a 4% PFA/PBS solution for 1 to 5 hours and washed with 1× PBS solution three times.

Kidneys were processed as follows. Mid-sections were sliced using a scalpel knife to obtain roughly 1- 1.5 mm thick transverse tissue slices containing all kidney regions from the renal cortex to the papilla. All procedures were performed in an incubator with an orbital shaker and temperature set to 37 °C. Either three channels (nuclei, carb, and amine) or two channels (carb and nuclei) were labeled using FLARE stains. For kidney tissues shown in **Fig. 1**, 3-channel tissue labeling, clearing, and imaging methods have been previously published (24). For kidney tissues in **Fig. 2-4**, kidney slabs were oxidized for 7 hours in 100mM NaIO_4_ in pH 5.0 NaOAc buffer, washed twice with 10 mL of NaOAc for 1 hour, and incubated overnight in 2 ml of 5 ug/mL AT565-hydrazide in 50% THF / 50% NaOAc buffer. For the reduction reaction, tissues were transferred to 100 mM NaCNBH_4_ in THF/NaOAc buffer for 1.5 hrs. Tissues were washed twice with 10 mL THF/NaOAc for an hour, and then labeled for nuclei by incubating overnight in 2 ml of 50% THF / 50% PBS with 10 µg/ml Sybr-Green I dye. Kidney tissues were incubated in a graded series of ethanol mixtures (25) with ascending ethanol concentration (5ml of 50%, 70%, 100%, and then 100% again) for 2 hours each at 37 °C. Tissues were placed in DCM for further delipidation until the tissue sank (∼10 minutes) and then transferred onto a glass vial containing ECi for 4 hours. ECi was refreshed once for a few hours before imaging in the same media.

Testes were processed as follows. Intact testes were delipidated in 50% CUBIC-L for 1 day and 100% CUBIC-L for 10 days. For carbohydrate labeling, tissues were oxidized for 5 hours in 100 mM NaIO_4_ in pH 5.0 NaOAc buffer, washed three times with 10 mL NaOAc for 30 minutes, and incubated overnight in 2 mL of 2.5 µg/mL AT647N-hydrazide in 50% THF/ 50% NaOAc buffer. For the reduction reaction, tissues were transferred to 100 mM NaCNBH_4_ in THF/NaOAc buffer for 1.5 hrs. Tissues were washed three times with 10 mL NaOAc for 30 minutes, and then labeled for nuclei by incubating overnight in 2 mL of 1.5 M NaCl in 1× PBS with 2.5 ug/ml propidium iodide, followed by one 30-minute wash in 1× PBS and two 30-minute washes in deionized water. Tissues were incubated in 50% CUBIC-R overnight and 100% CUBIC-R for 2 days for refractive index matching before imaging.

### Imaging with a custom open-top light-sheet (OTLS) microscope

High-resolution volumetric images of ECi-cleared mouse kidney slices were rapidly acquired (15 minutes/mm^3^) using OTLS microscopy. Data used for validating the performance of TubuleMAP (**Fig. 2c-f; Supplementary Fig. S3**) were captured with a previously published, custom-built OTLS microscope (45) with sub-micron lateral and 3.5 μm axial resolution. Data for kidney and testis studies (**Fig. 2, 3, 4, 5**) were captured with a redesigned multi-resolution OTLS microscope based on (17). This system incorporated a new four-channel (405, 488, 561, and 638 nm) laser package (Versalase, Vortran Lasers) for illumination. Light from the laser was shaped into a pencil beam that was scanned across the field of view using two-axis galvanometers (6220H, Cambridge Technology) to generate the light sheet. Fluorescence emission was collected via a FOV-maximized optical path as described in Glaser et al. (17) and focused onto a 2048 × 2048 pixel sCMOS camera (ORCA-Flash4.0 V3, Hamamatsu) at a sampling of ∼0.42 μm per pixel. Using gold nanoparticles, the spatial resolution was determined to be ∼1 μm lateral and 2.9 μm axial with a refractive index of 1.56. The exposure time per frame was set to 20 ms. The specimen was laterally scanned across the lightsheet to generate image strips with 5% tile overlap. When tiling vertically, the laser power was increased with depth to account for laser attenuation as it penetrates deeper into the specimen. 3D imaging was achieved using a combination of stage scanning and lateral/vertical tiling with a motorized XY stage and Z actuators (S562-2235A and LS-50-FTP, Applied Scientific Instrumentation). A TTL SYNC trigger signal was sent from the XY stage to a DAQ card (PCIe-6738, National Instruments), which in turn sent synchronized output voltages to the camera, lasers, and galvos. The camera was operated in synchronized readout mode to ensure reproducible exposure for each image strip. The lateral scanning mirror was actuated with a sawtooth waveform that completed a single period within the total exposure time of the raw camera frame. The entire image acquisition was controlled by a custom-written Python program available from the authors upon request.

During acquisition, the images were collected by a dedicated custom workstation (Puget Systems) equipped with a high-specification motherboard (Asus WS C422 SAGE/10 G), processor (Intel Xeon W-2145 3.7 GHz 8 Core 11MB 140 W), and 256 GB of RAM. The motherboard houses several PCIe cards, including CameraLink frame grabbers (mEIV AD4/VD4, Silicon Software) for streaming images from the camera, a DAQ card, and a GPU (TitanXP, NVIDIA). Datasets were streamed to a local 4 TB M.2 NVMe drive (Samsung) capable of outpacing the data rates of the microscope system. Data were transferred to an 80 TB RAID array storage drive for backup.

### Imaging data post-processing

Raw camera frames were saved in real-time in a hierarchical data format version 5 (HDF5) file, with on-the-fly downsampling (×2, ×4, ×8, and ×16) applied along all axes to generate a resolution pyramid. Simultaneously, GPU-based B3D (46) compression (5-10×, signal-preserving) was applied to selectively reduce image noise and minimize file size while preserving image features. The HDF5 file and metadata were imported into BigStitcher (47) (ImageJ plugin), where imaged tiles were aligned, stitched, and fused at the highest resolution into one contiguous 3D volume. The resulting dataset was exported as a chunked, multiscale OME-Zarr (26) file for downstream analysis. The OME-Zarr format enables efficient random access and straightforward parallelization, allowing us to run multiple tracking processes in parallel; one process per tubule.

### Methods description for TubuleMAP software

All algorithmic components described below were implemented in the open-source TubuleMAP software (available at https://github.com/Davidrbr95/TubuleMAP/tree/main), which contains the full analytical workflow used for all reported results. The descriptions provided in this manuscript and supplementary materials summarize the main algorithmic steps, while the repository provides the exact implementation.

### Overview of TubuleMAP reconstruction process

Starting from two user-defined seed points, TubuleMAP performs stepwise vector tracking to follow the tubule centerline. At each step, the forward direction is estimated from the two most recent centerline nodes, and the next node location is proposed using an adaptive step size **(Supplementary Fig. S2a**).

A 2D plane orthogonal to the forward direction is then sampled at the proposed location and segmented to identify the tubule cross-section (**Supplementary Fig. S2b**). If multiple candidate masks are detected, the mask overlapping the plane center is selected. The proposed node is then repositioned to the centroid of the selected mask, and this correction is used to update the forward direction for the next iteration. Repeating this procedure yields a sequence of centered orthogonal cross-sections along the tubule path, which are interpolated in 3D and voxelized at native image resolution to generate a 3D pixel-level reconstruction.

To improve robustness across changes in tubule diameter, morphology, and staining, TubuleMAP uses three recovery modules that are invoked only when needed to save time: adaptive parameter scaling, rotational search, and troubleshooting and model switching **(Supplementary Fig. S2c-f**). Together, these modules adjust tracking parameters, correct local orientation errors, and recover cross-sections when standard segmentation fails. If these automated procedures still do not produce a suitable mask, the track is flagged for correction in a Napari-based human-in-the-loop interface and then returned to the processing queue.

### Adaptive tracking parameters (AP)

At each step, we compute three runtime parameters from the most recently segmented cross-sections: the node step size (*S*), the cross-section plane dimensions (*D*), and the segmentation diameter jitter (*j*; see Troubleshooting and model switching). From the segmented cross-section, we obtain area (*A*) and calculate its equivalent circular diameter 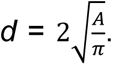 To reduce frame-to-frame noise, the running average (*d*^′^) is computed over the previous *W* and clamped to user-provided physiological bounds [ *d_min_*, *d_min_*]. The runtime parameters are then derived from *d’* using user-set scaling factors: *S = round(d’/k_step)*, *J = round(d’/k_jitter)*, and *D = round(d’*k_dim)*. The diameter was initialized at 34 μ*W* ￼*d_min_*, *d_max_*o [￼] = [0.8 μm, 50 μm]. The scaling factors were *k_jitter* = 3, *k_step* = 5, and *k_dim* = 2.5.

### Rotational search (RS)

Rotational search was performed only when the segmented cross-section was eccentric, indicating a possible bend or local orientation error; otherwise, this step was skipped to save time. The goal of the rotation module is to maintain accurate, stable reconstruction through tight bends. After segmenting the sampled plane, we fit an ellipse to the cross-section and calculate its in-plane orientation *φ*. With the frame’s axes (*u, v*), we define a rotation axis aligned with the ellipse’s minor axis (*r = -sin φ u + cos φ v*). We generate a small set of candidate planes by rotating about *r* by few degrees (−60° to 60° with a step size of 10°) and re-segment each plane to obtain candidate cross-sections. The candidate plane yielding the lowest eccentricity (roundest cross-section) is selected; its mask becomes the final cross-section for volumetric reconstruction, and its normal becomes the new forward direction.

### Troubleshooting and model switching (TS)

When segmentation fails to generate a mask, TubuleMAP invokes a two-stage troubleshooting workflow. In stage 1, the current segmentation model is applied to a small local volume over a diameter sweep using (*d’*), (*d’ - J*), and (*d’ + J*), yielding three candidate segmentation volumes (**Supplementary Fig. S2e**). The local volume is centered on the target plane, aligned with the forward tracking direction, and extends to a depth of 10 slices with the target plane as the first slice. These competing hypotheses are integrated with Ultrack (27), which generates a consensus segmentation that accounts for depth consistency. If successful, the center-overlapping mask from the consensus segmentation on the target plane is used to continue the reconstruction. If stage 1 fails to generate a mask, stage 2 repeats the procedure (diameter sweep and Ultrack integration) across the full segmentation-model suite, including models specialized for distinct tubule morphologies and staining patterns. The final center-overlapping masks on the target planes of the Ultrack consensus and each model–diameter combination are compared, and the model with the highest intersection over union (IoU) is used in subsequent iterations (**Supplementary Fig. S2f**).

### Cellpose model suite for nephron reconstruction

We trained a suite of Cellpose-based models for nephron reconstruction, including segment-specific models for the PCT, thin limb, and thick limb, to support model switching during inference. Training data were derived from 5 manually curated centerlines (held out from validation) that were uniformly resampled to produce evenly spaced orthogonal planes centered on the nephron. Plane dimensions were set by morphology (PCT: 168 × 168 μm^2^; thin limbs: 84 × 84 μm^2^; TAL: 168 × 168 μm^2^) to satisfy Cellpose’s training requirements (approximately five masks per image). Since Cellpose models trained on ∼500-1000 masks show robust performance (22), we selected ∼100 diverse planes per segment using multi-scale structural similarity index measure (SSIM); a frame was kept only when it met both local (*α*) and global (*β*) change criteria and was at a minimum interval (*t*; measured in pixels) from a previously kept plane (PCT: *α*=0.08, *β*=0.08, *t*=20; thin limb: *α*=0.3, *β*=0.08, *t*=5; thick limb: *α*=0.5, *β*=0.2, *t*=10). To generate labeled data, each plane was segmented by Cellpose’s cyto2 model and then manually corrected by an expert annotator. Three specialized segmentation models were subsequently trained on corrected datasets.

### Tracking error quantification

Overall tracking performance was evaluated on a set of 20 nephrons with manually annotated centerlines (**Fig. 2c**), using breaks per nephron as the primary metric. A break was defined as any point at which tracking could no longer correctly follow the manual path, including excessive deviation from the ground-truth centerline, backtracking, or failure to generate a segmentation mask. After each break, tracking was automatically restarted from the point of deviation. To benchmark overall tracking performance, we compared three variants of TubuleMAP under identical seeds, stopping rules, and initializations: TubuleMAP without AP, RS, or TS (pure vector-based tracking) using only Cellpose cyto2; TubuleMAP with AP, RS, and TS enabled using only Cellpose cyto2; and TubuleMAP with AP, RS, and TS enabled using our segment-specific model suite. To isolate the contribution of each module (AP, RS, and TS), we also evaluated all eight on/off permutations of AP, RS, and TS with and without the model suite (**Supplementary Fig. S4a**). **Supplementary Fig. S4b** presents the same ablation study using uninterrupted tracking distance.

### Timing experiments

The pipeline scalability was evaluated by reconstructing multiple instances of a randomly selected nephron in parallel, increasing the number of concurrent processes. On an Ubuntu 22.04 workstation (GPU: RTX 4090; 270 GB RAM), runs scaled up to 30 instances; beyond this, GPU memory was exhausted, and processes began to fail. Each run was timed individually, and the average run-time per number of concurrent processes is reported (**Supplementary Fig. S4e**).

### Evaluation of reconstruction quality

Two nephrons were manually segmented in ImageJ by annotating consecutive 2D slices. On average, manual annotation required ∼40 hours per nephron. These manual annotations were compared with the corresponding TubuleMAP reconstructions using intersection over union (IoU), F1 score, precision, and recall (**Supplementary Fig. S5**).

### Post-tracking analysis

After tracking was complete, parameters like length, curvature, torsion, and diameter for individual trajectories were calculated using custom Python scripts. Straightened views were generated from the Zarr file volume as an HDF5 image stack for each tubule, and the GUI was configured to also allow for manual corrections to the 2D segmentations per plane, if desired. Complete 3D reconstructions were rendered in Blender v4.2.

### Nephron Cytometry

For the nuclei quantification, a bounding box was generated from each nephron mesh and the same region was cropped from the nuclei channel dataset. 3D nuclei segmentation was performed on the cropped volume using Cellpose pretrained ‘nuclei’ model and the centroid of each nucleus was obtained. Nuclei were assigned to the nephron by selecting nuclei whose centroid was located within the nephron mesh. After nuclei assignment, nuclei density per unit length and per unit volume were calculated using consecutive 20 µm windows along the nephron centerline. Nephron segment-level values were reported as the mean across all windows within the corresponding segment. Nuclei meshes were generated for visualization in Meshlab software.

### nnU-Net model training for tissue regions framework

An nnU-Net model was trained to generate 2D segmentation masks corresponding to the four tissue regions (cortex, OSOM, ISOM, and IM regions) per slice of the kidney volume. nnU-Net provides an end-to-end semantic segmentation workflow that adapts its pipeline to a dataset and includes automated pre- and post-processing. 5 image slices spanning the volume and containing 4-class segmentation masks were used as a training dataset while 5 other slices were used as a test dataset. The resulting model was used to predict 4-class segmentation for the remaining ∼2900 slices in the volume.

### Reconstruction and analysis of seminiferous tubules

Seminiferous tubules were reconstructed with TubuleMAP using the Cellpose ‘cyto3’ model applied to the carbohydrate channel. Unlike nephron reconstructions, which required specialized segmentation models, seminiferous tubules could be segmented adequately with this general-purpose model because the staining was more homogeneous and well-defined. While nephrons provide clearer anatomical landmarks (e.g., the glomerulus) for the placement of user-defined seed points, seminiferous tubules form a branching network with less obvious start and end points. Tracking was therefore initialized from randomly selected seed points across tubules and continued until a track terminated in the rete testis or merged with a previously reconstructed tubule. Overlaps were identified using Euclidean distance between the centerlines, after which one reconstruction was terminated and merged with the other. Branch points were identified as areas where 3 or more reconstructed segments were merged. Reconstruction continued until visual inspection indicated that all seminiferous tubules had been completely reconstructed.

After reconstruction, seminiferous tubule cross-sections were extracted and classified using an automated hierarchical classifier into three visually distinct classes based on DNA staining patterns: the dense clusters class corresponds to stages I–V and X–XII, containing bright spermatozoa lining the rim of a thick epithelium; the sparse clusters class corresponds to stages VI–VII, containing bright, elongated spermatids forming a thin epithelium; and the dark class corresponds to stages VIII–IX, containing bright spermatozoa lining the rim of a thick epithelium. Frames were classified as dark when the spermatozoa area identified by a simple intensity threshold with parameter *a* occupied less than a fraction *b* of the tubule cross-sectional area. The remaining frames were re-segmented with a second intensity threshold *c* and classified to the sparse clusters class if the two largest clusters identified by connected component analysis were below the threshold *d*; else frames were assigned to the dense clusters class. Parameters were optimized via grid search on manually labeled cross-sections from two seminiferous tubules sampled every 20 pixels (training = 273). Performance was then evaluated on a separate manually labeled tubule processed in the same way (testing *n* = 373). The optimal parameters were *a* = 2.5, *b* = 0.02, *c* = 86, and *d* = 0.62, yielding an F1 score of 0.86 (**Supplementary Fig. S10c**). Because the automated classifier occasionally produced frame-to-frame fluctuations, class labels were smoothed along the tubule axis using a 2-mm rolling-window, producing the final classifications that closely matched the visually distinct patterns observed in straightened tubule views (**Supplementary Fig. S10d**).

## Data Availability

A subset of the multi-terabyte datasets and trained segmentation models can be made available on reasonable request from the authors.

## Code Availability

The repository for the TubuleMAP source code and its updated versions are available at https://github.com/Davidrbr95/TubuleMAP/tree/main.

## Acknowledgements

The authors thank Dr. Adam Glaser and Dr. Chenyi Mao for providing access to a published mouse-kidney dataset used in the paper for benchmarking TubuleMAP. The authors also thank Hung-Yu Chang, Hoyin Lai, and Chi-Chou Huang at Aivia, Leica Microsystems, for early advice and support on various aspects of this work. This work was funded in part by the NIH grants R21 DK136026 (J.C.V.), R01 DK135716 (S.J.S., J.C.V.), R01 CA244170 (J.T.C.L.), R01 EB031002 (J.T.C.L.), R01 CA268207 (J.T.C.L.), R01 DK138948 (J.T.C.L.), TL1DK143270 (P.V.). Indiana O’Brien U54 DK137328 (P.C.D., J.C.V, and J.T.C.L.), NIDDK Innovative Science Accelerator Program ISAC DK128851 (C.P.) and Washington Research Foundation postdoctoral fellowships (C.P. and D.B.).

## Contributions

C.P., D.B., W.X., and J.C.V. designed the studies. C.P. built the open-top lightsheet microscope where most data were acquired. C.P., W.X., R.M.S, M.M.M., and D.G.E. prepared the mouse kidney tissues. C.P. imaged the kidney tissues. Y.G. and P.V. prepared and imaged the mouse testis samples. C.P., D.B., and W.X. trained the Cellpose models and wrote the tracking algorithm. C.P., W.X., Y.G., P.V., N.M., E.K., N.F., Z.G., and N.J.P. performed manual annotations and corrections. J.T.C.L. and S.J.S. provided partial funding and supervisory support. J.C.V. supervised and guided the entire project. All authors prepared the paper.

## Corresponding authors

Correspondence to Chetan Poudel or Joshua C. Vaughan.

## Supplementary Information

### Supplementary Figures

**Supplementary Figure S 1:**
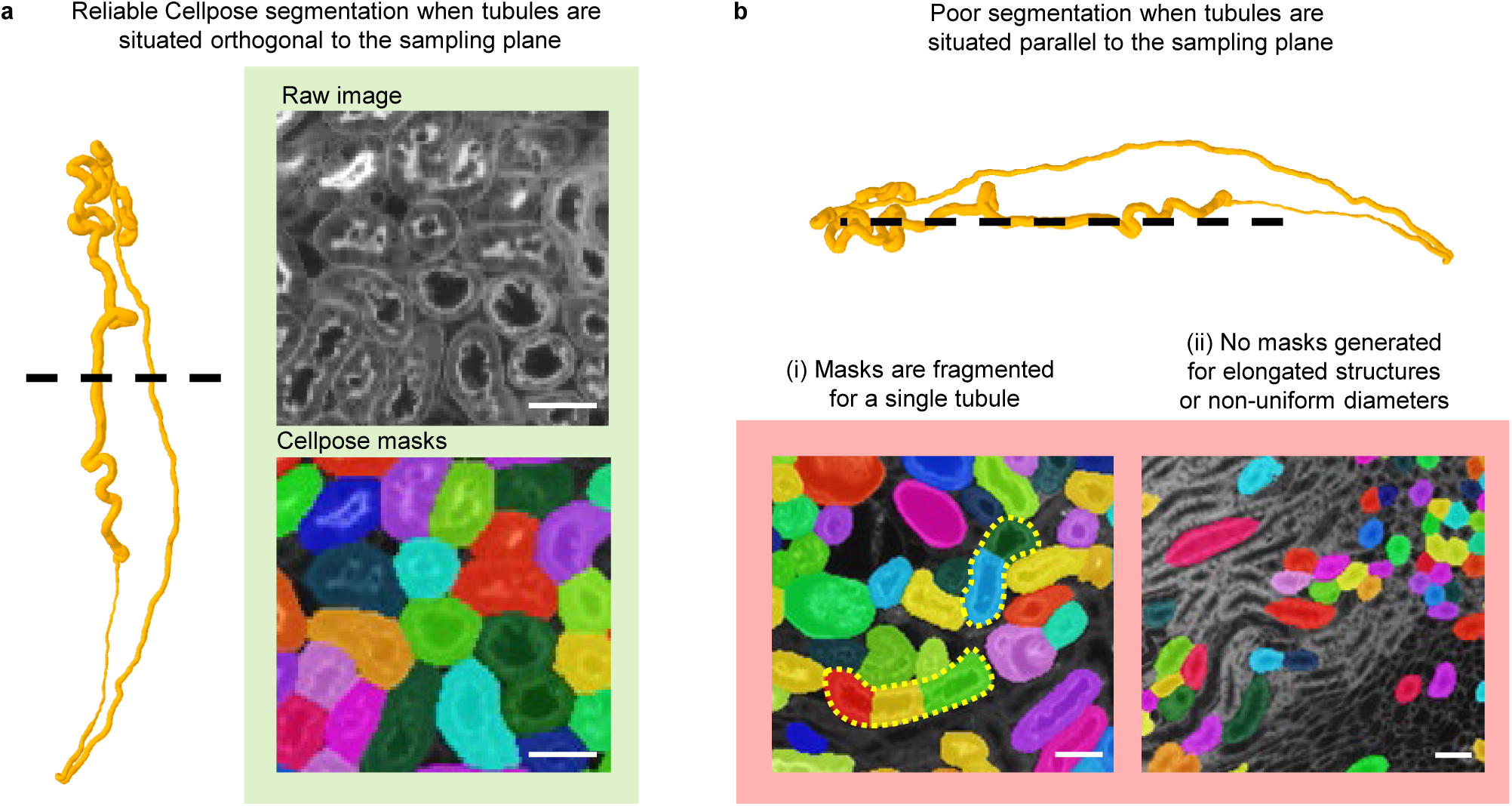
Segmentation performance varies with spatial orientation of tubules. (**a**) Cellpose segmentation shows reliable performance when tubules are situated orthogonal to the sampling plane, cross-sections are mostly circular, and diameters are uniform. (**b**) Cellpose segmentation is poor when tubules are situated parallel to the sampling plane, yielding elongated cross-sections. (i) A single tubule cross-section generates several fragmented segmentation masks (e.g. yellow dashed lines) that need manual joining/correcting, or (ii) most tubule cross-sections do not generate a mask due to elongated structures or different diameters.

**Supplementary Figure S2:**
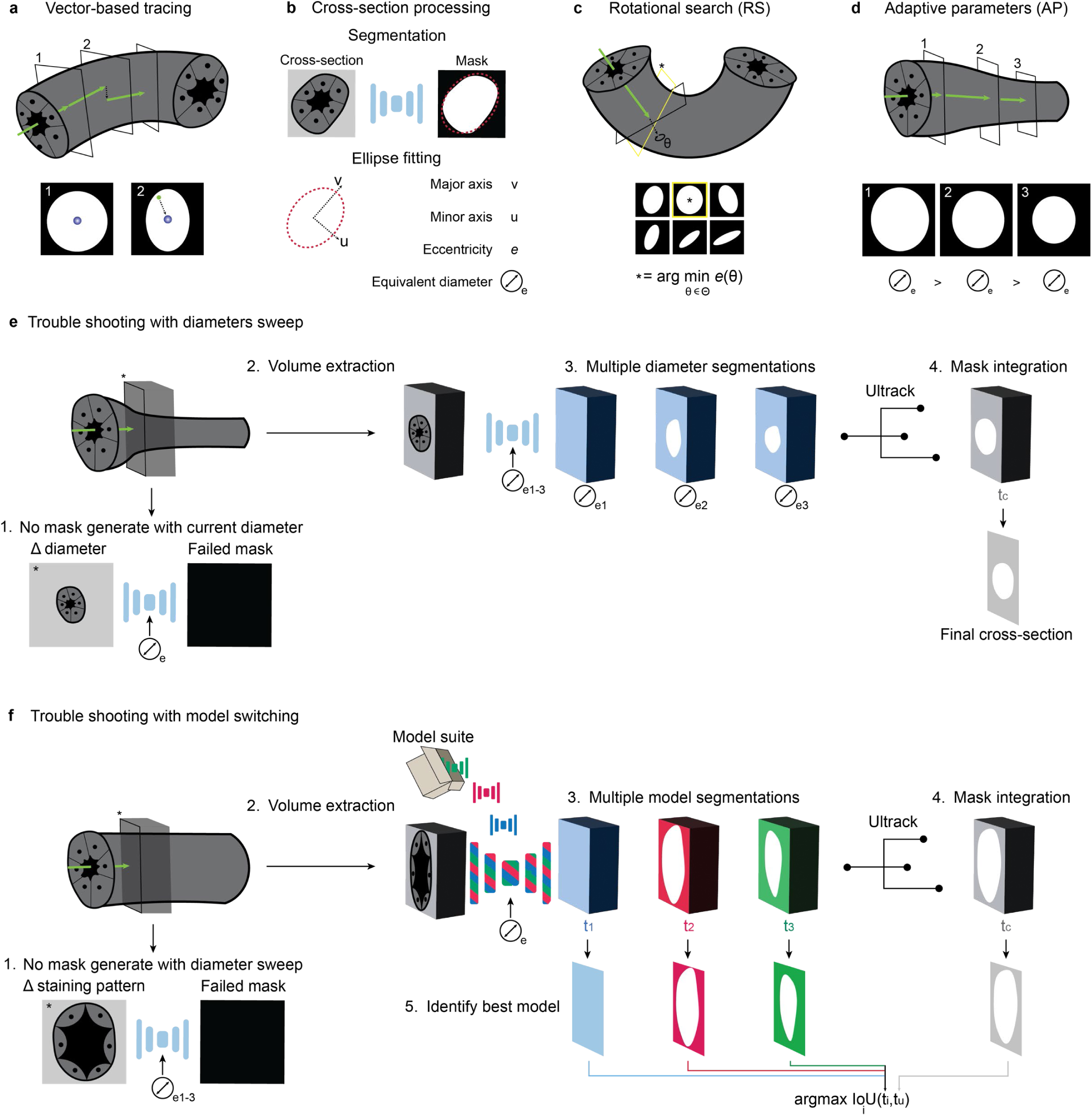
TubuleMAP modules. (**a**) Cross-sections obtained at each step are segmented to obtain a mask, which is fit to an ellipse shown by the red dashed outline to quantify its centroid, eccentricity, and equivalent diameter. (**b**) The tracking vector from the previous step is first used to predict the direction for the next step, but is adjusted after segmentation towards the mask centroid. (**c**) At tubule bends where the mask eccentricity can exceed a set threshold, a rotational search is invoked to select the direction with the minimum eccentricity most circular cross section). (**d**) The equivalent diameter obtained from the latest mask is used to update parameters and adapt to tubule size changes. (**e**) If segmentation fails, a diameter sweep is performed on a local volume sampled along the vector and an optimal diameter is chosen by maximizing overlap with the aggregated mask set. (**f**) If segmentation fails on a diameter sweep, all available segmentation models are applied on the same sampled volume and an optimal model is selected by the same overlap metric. For future steps, we revert to the simpler vector-based tracking using the new optimal model and additional modules are invoked as needed.

**Supplementary Figure S3:**
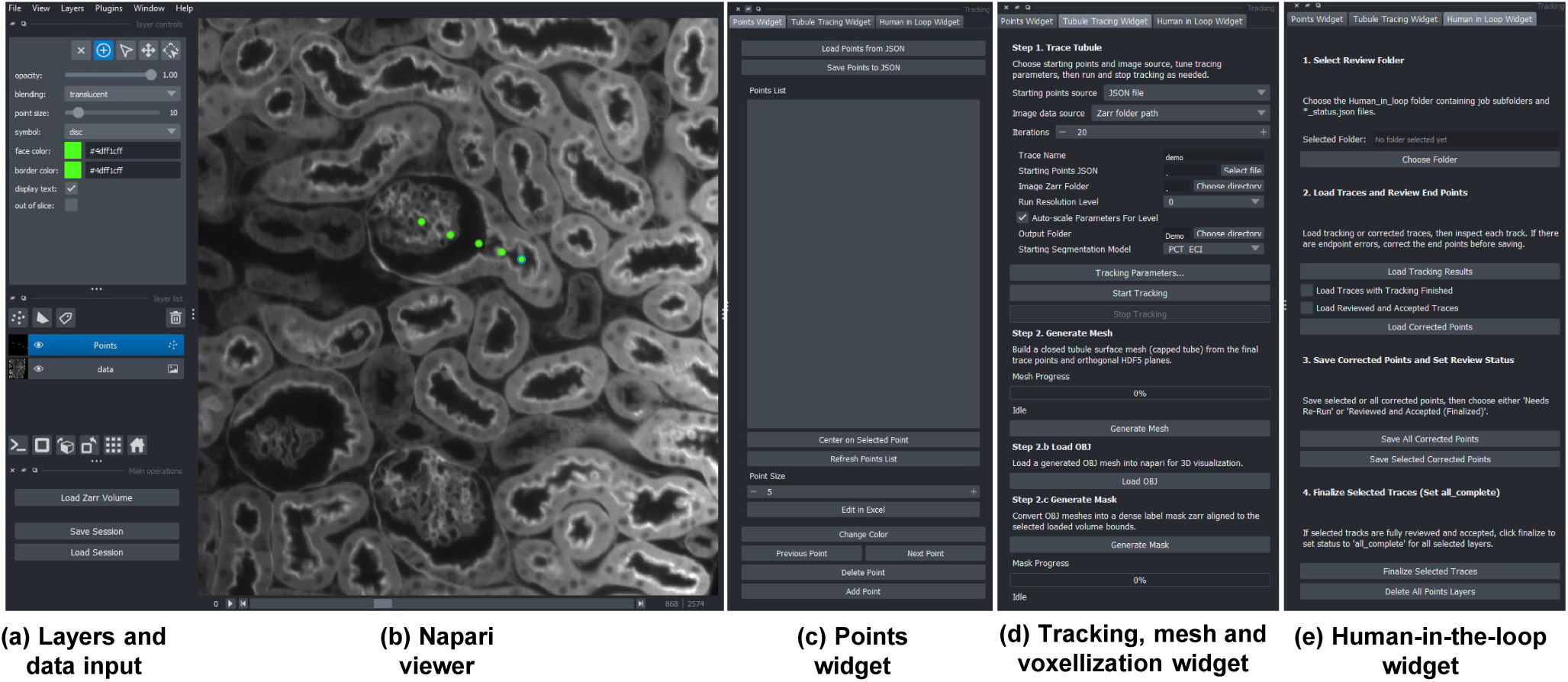
Screenshot of the TubuleMAP interface and associated features. (**a**) Widget for layers and data input allows for loading zarr and tif files and creating new points/annotation layers. (**b**) The napari viewer displays the image and provides an interface for human intervention. (**c**) The points widget enables centering of the image on a selected point, toggling between previous and next points, adding or deleting points, and changing colors of individual tracks for visualization. (**d**) The tracking widget lets a user choose various tracking parameters to perform a single tracking run on the GUI itself. The output trajectory is displayed immediately on the napari viewer after each run. 3D meshes based on the segmentation can be generated and loaded directly here, and they can also be voxellized to generate full 3D masks. For high-throughput tracking, a separate multiprocessing script is used and many output tracks can be displayed simultaneously on the GUI. (**e**) The human-in-the-loop widget allows for batch loading and saving tracks with specified status (‘running’, ‘needs correction’, ‘corrected’, or ‘complete’) to quickly review and to organize many files.

**Supplementary Figure S 4:**
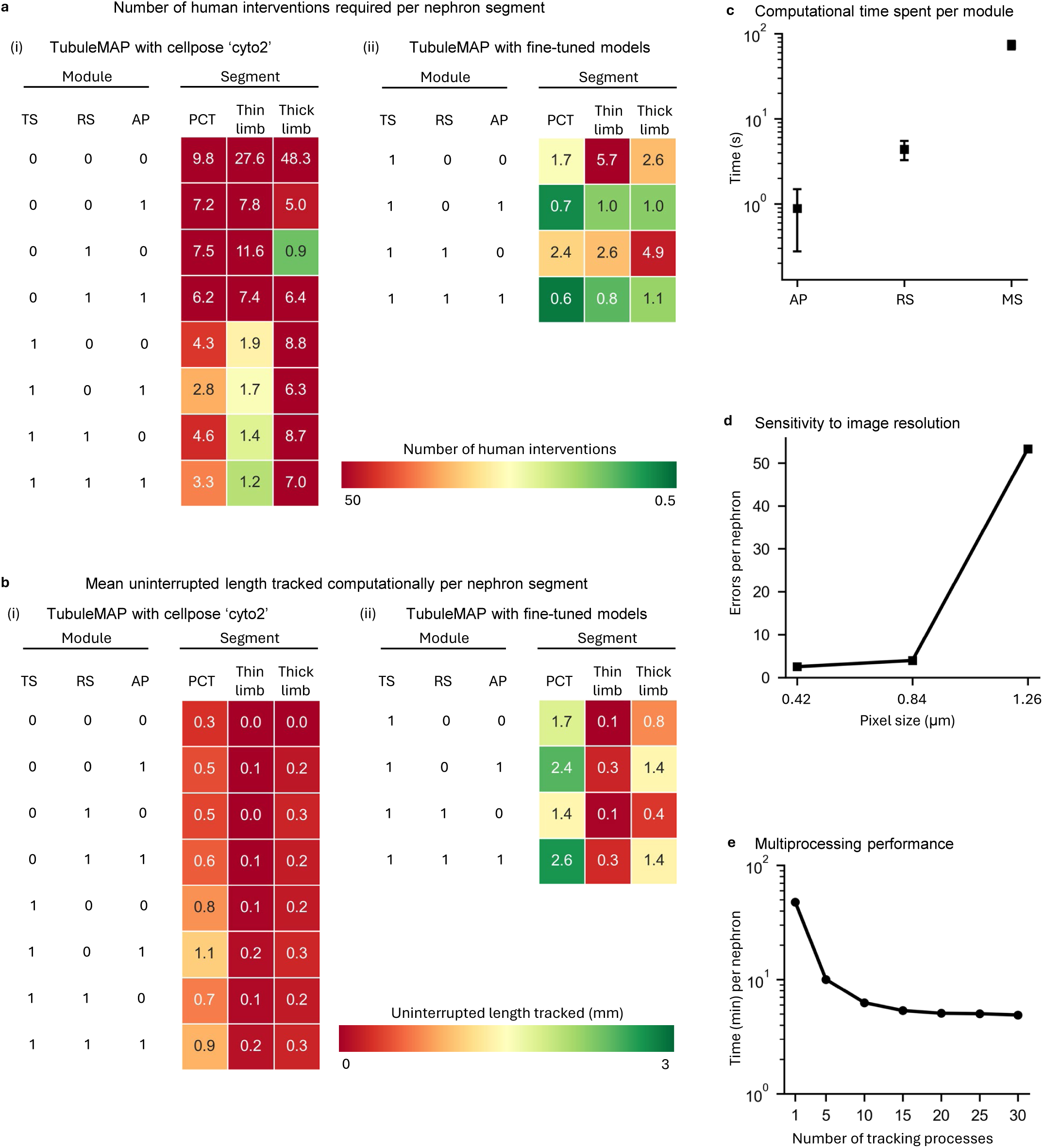
Characterization of TubuleMAP performance, speed, and scalability. (**a**) Ablation studies characterizing the number of human interventions required for generalist (Cellpose ‘cyto2’) vs fine-tuned models for tracking different nephron segments after inclusion of various TubuleMAP modules. MS: model switching, RS: rotational search, AP: Adaptive parameters, PCT: proximal convoluted tubule. (**b**) Mean uninterrupted tracked length per nephron segment across TubuleMAP configurations for generalist and fine-uned models. (**c**) Speed: compute time per step for each TubuleMAP module. (**d**) Resolution sensitivity: impact of spatial resolution on tracking performance. (**e**) Scalability: reduction in tracking time per nephron after parallel rocessing of tracks on a single NVIDIA RTX 4090 GPU.

**Supplementary Figure S 5:**
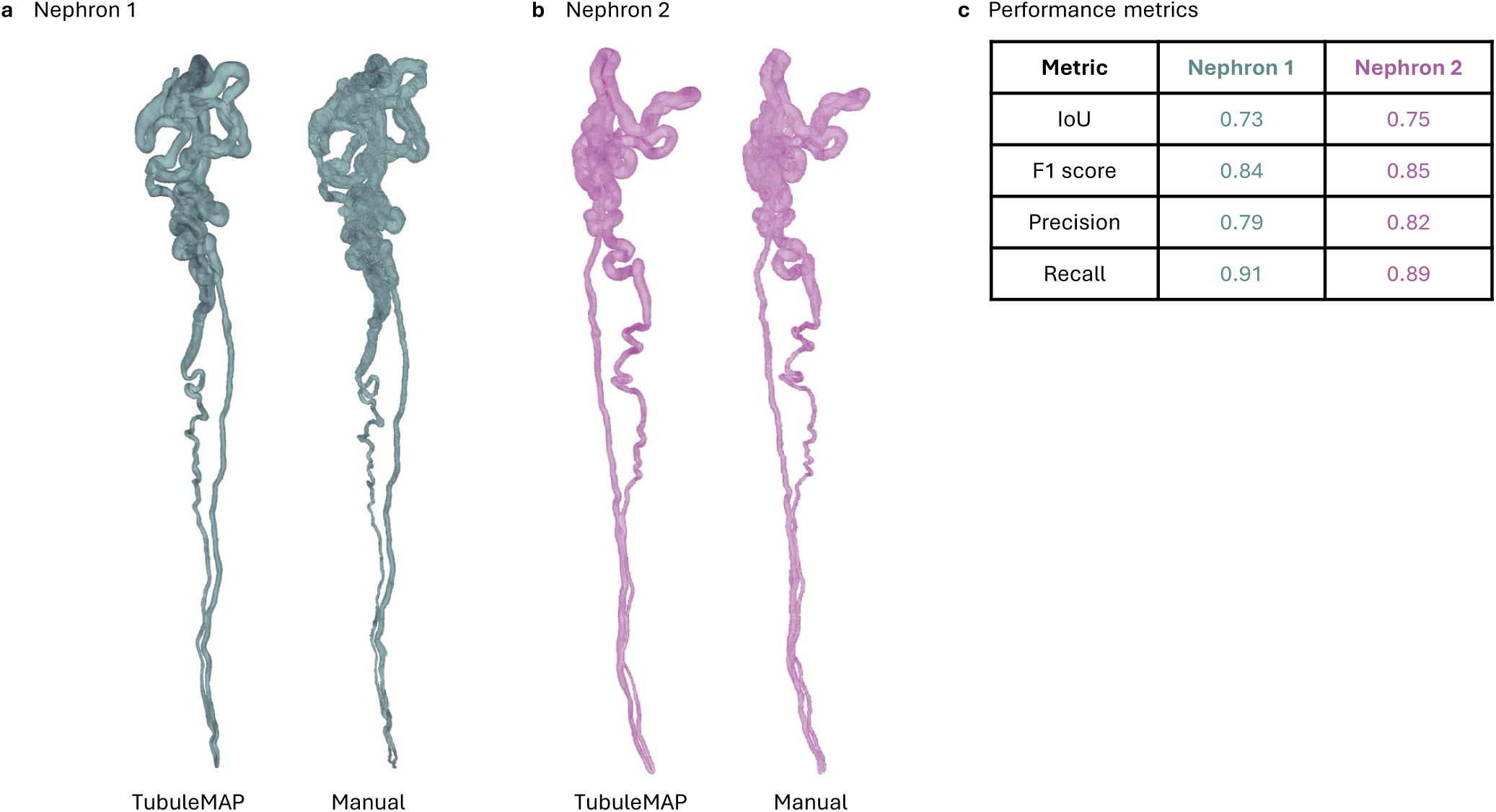
Comparing 3D segmentation quality between a manual approach versus TubuleMAP for two nephrons. (**a-b**) 3D mesh reconstructions of two nephrons generated by TubuleMAP and manual annotation. Nephron 1 is shown in (**a**) and Nephron to shown in (**b**). (**c**) Quantitative performance metrics for each nephron, including Intersection over Union (IoU), Dice score, precision, and recall.

**Supplementary Figure S 6:**
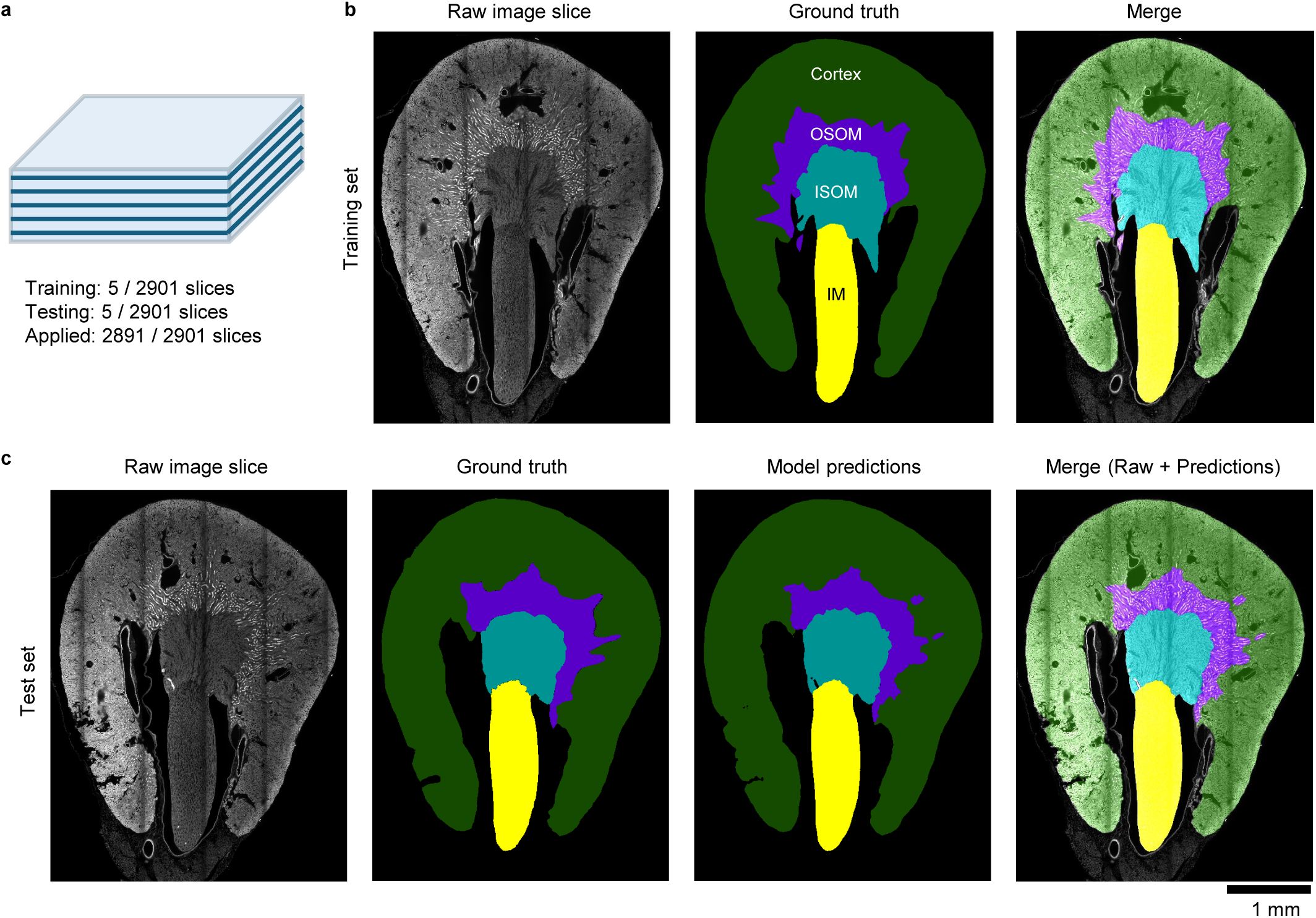
nnU-Net based segmentation of kidney tissue regions. An nnU-Net model, designed to handle diverse biomedical imaging datasets, was trained to segment four kidney tissue regions: cortex, outer stripe of outer medulla (OSOM), inner stripe of outer medulla (ISOM), and inner medulla (IM). (**a**) Ten slices spanning the whole volume were manually annotated: five slices (raw images and corresponding ground-truth masks) were used for training and five for testing. (**b**) An example slice (raw image, corresponding ground-truth masks, and merge) used for training is shown. The model achieved high intersection-over-union (IoU) scores across classes: (IM, 0.99; ISOM, 0.93; OSOM, 0.84; cortex, 0.96), with a mean IoU of 0.93. (**c**) An example slice from the test set comparing ground-truth masks and model predictions is also shown. The trained model was subsequently applied to the remaining slices to generate a 4-class segmentation of the entire volume.

**Supplementary Figure S 7:**
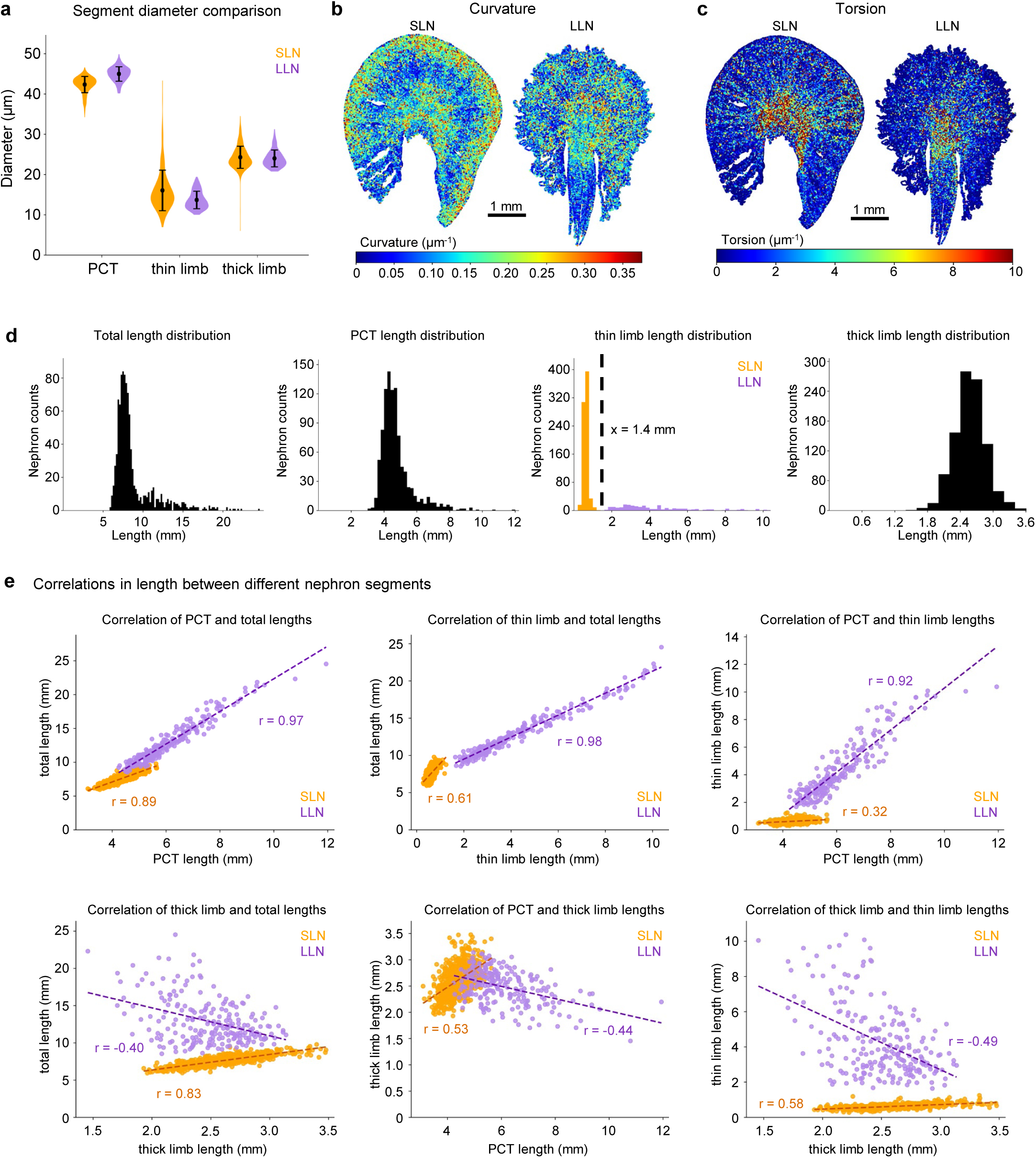
Quantification of mesoscale organization of the kidney. (**a**) Violin plots of segment diameters for SLN and LLN nephrons (PCT, thin limb, and thick limb). Mean ± S.D. is shown. (**b-c**) Mesoscale distribution of curvature (b) and torsion (c) computed from all nephron traces. Values were downsampled by taking the maximum absolute value within each 20 × 20 µm^2^ spatial bin in a 2D projection. (**d**) Histograms of total nephron length and segment-specific lengths (PCT, thin limb and thick limb) with a bin size of 0.2 mm. Thin limb lengths show a bimodal distribution and a length of 1.4 mm was used as the threshold to classify nephrons as SLN or LLN. (e) Pairwise correlation between lengths of different nephron segments and the total nephron length. r is Pearson correlation coefficient.

**Supplementary Figure S8:**
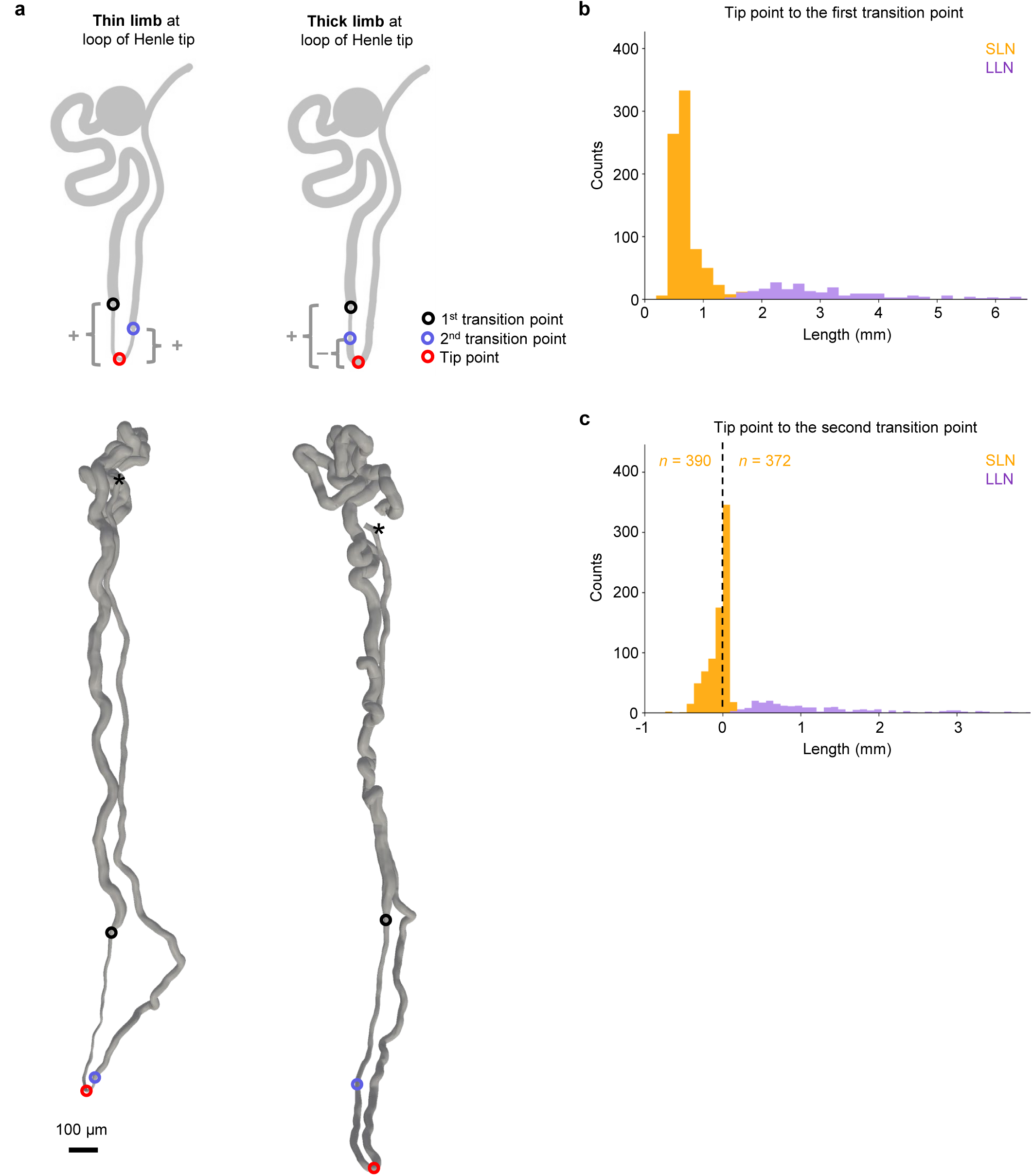
Characterizing the presence of thin limb or thick limb at the loop of Henle tip. (**a**) Schematics and representative reconstructed nephron meshes showing either the thin limb or the thick limb at the loop of Henle. The first transition point corresponds to the PCT to thin limb transition. The second transition point corresponds to the thin limb to thick limb transition. The sign convention shown in (a) is applied in (b-c). (**b**) Histograms of distances from the tip point to the first transition point for SLN and LLN. The first transition is always before the tip point (all positive values) (**c**) Histograms of distances from the tip point to the second transition point for SLN and LLN. For LLN, all second transitions are after the tip point (all positive values). For SLN, the second transition point can be before or after the tip point (negative and positive values).

**Supplementary Figure S9:**
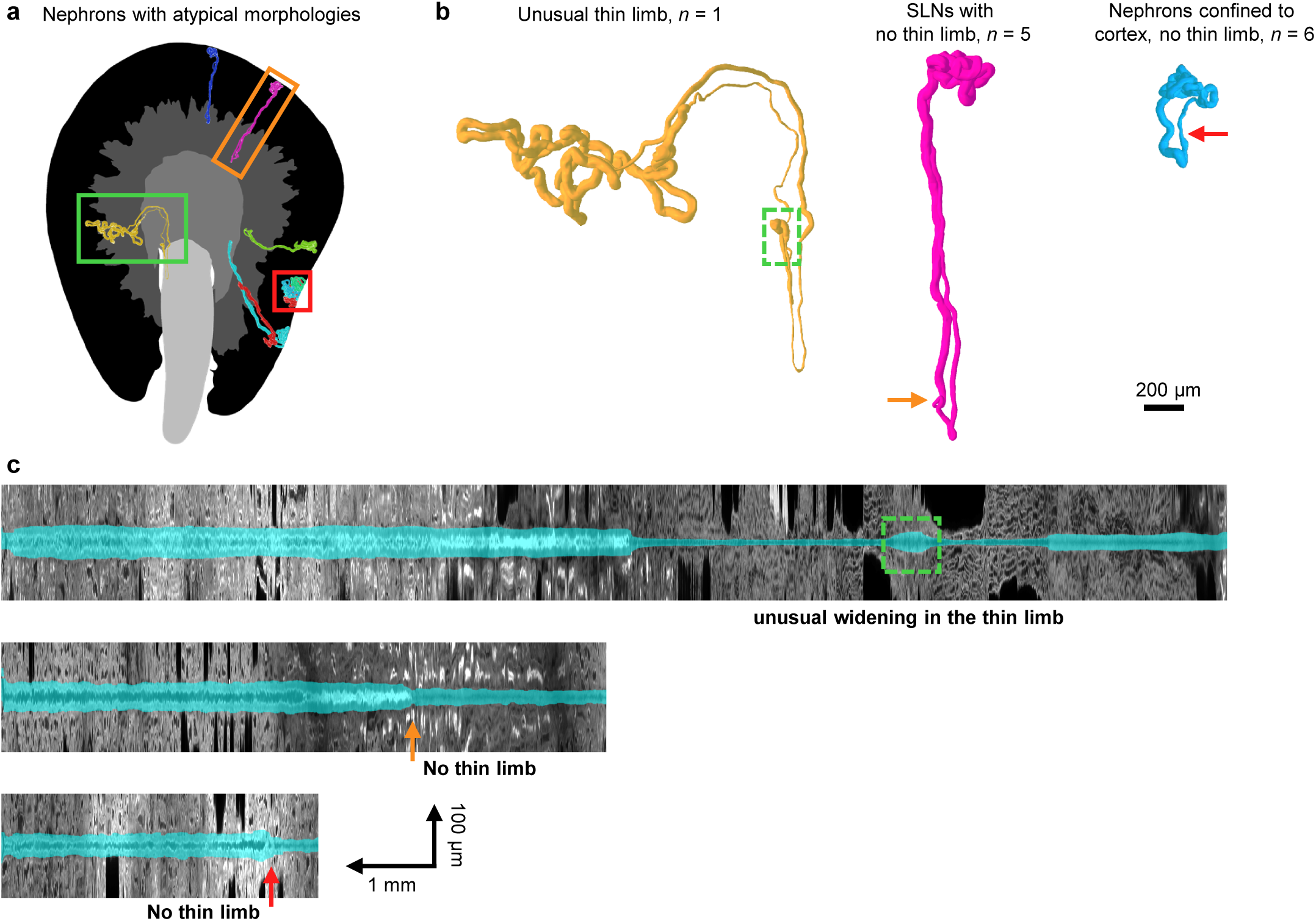
Nephrons with atypical morphologies. (**a**) Spatial localization of nephrons showing atypical morphologies within corresponding tissue regions. Colored boxes mark the examples shown in (b). (**b**) Three representative mesh reconstructions illustrating the classes of atypical morphologies: a nephron with unusual widening within the thin limb (*n* = 1), SLN with no detectable thin limbs (*n* = 5) and nephrons confined to cortex without detectable thin limb (*n* = 6). (**c**) Representative straightened views with segmentation overlay. The green dashed box highlights an unusual widening that appears like the thick limb between thin limbs. Orange and red arrows indicate the transition point from PCT to thick limb.

**Supplementary Figure S 10:**
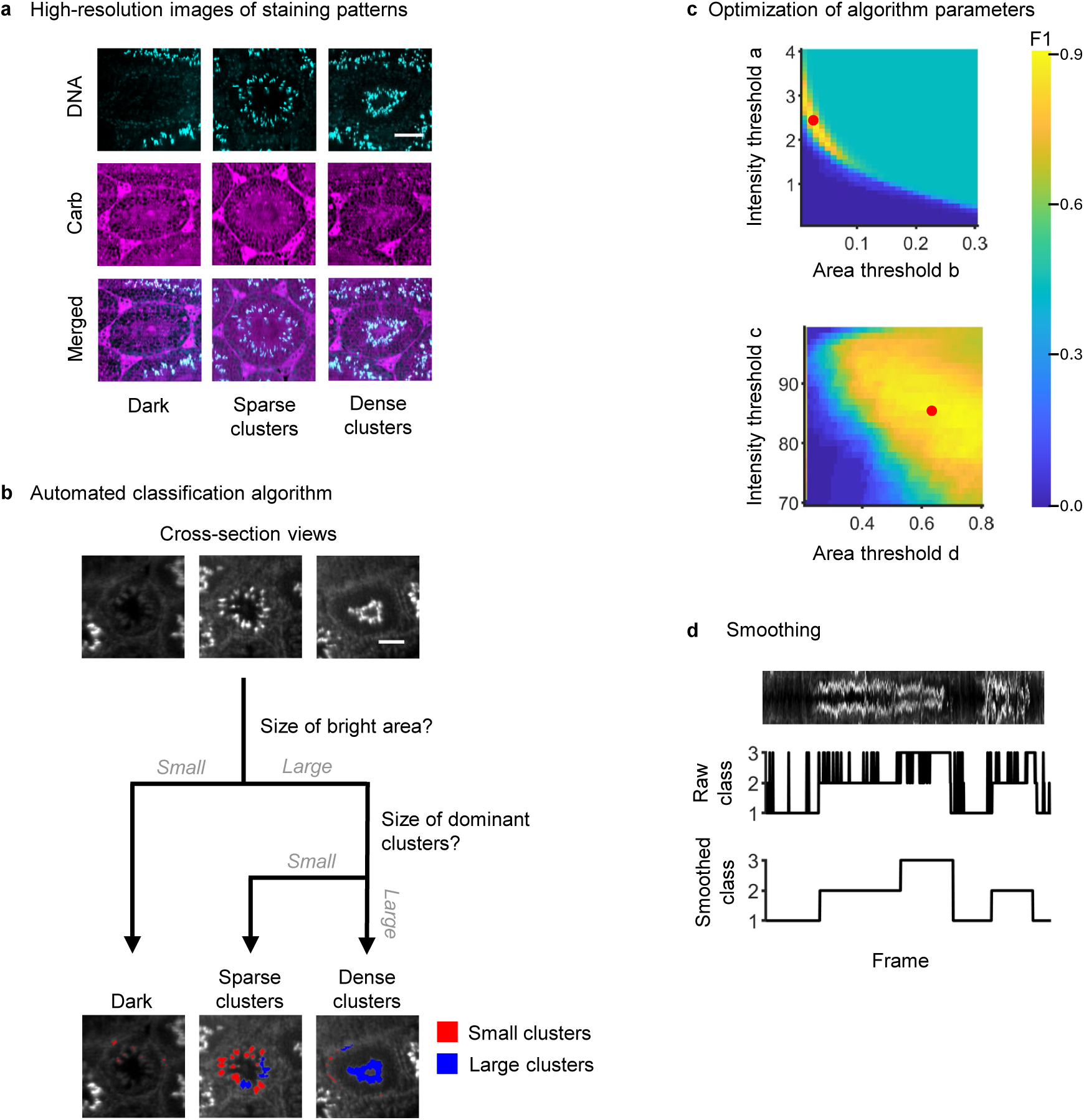
Analysis of spermatogenic waves in seminiferous tubules. **(a**) High-resolution images of carbohydrate and DNA channels exemplifying the three assigned classes: dark, sparse clusters, and dense clusters. (**b**) Schematic for hierarchical classification of seminiferous tubule cross-sections into the three classes using the DNA channel. (**c**) F1 score across the parameter optimization space. The optimal operating point is shown in red. All scale bars are 75 µm. (**d**) Straightened view of a seminiferous tubule, shown together with the raw classification and the smoothed classification obtained by a rolling mode filter.

**Supplementary Figure S11:**
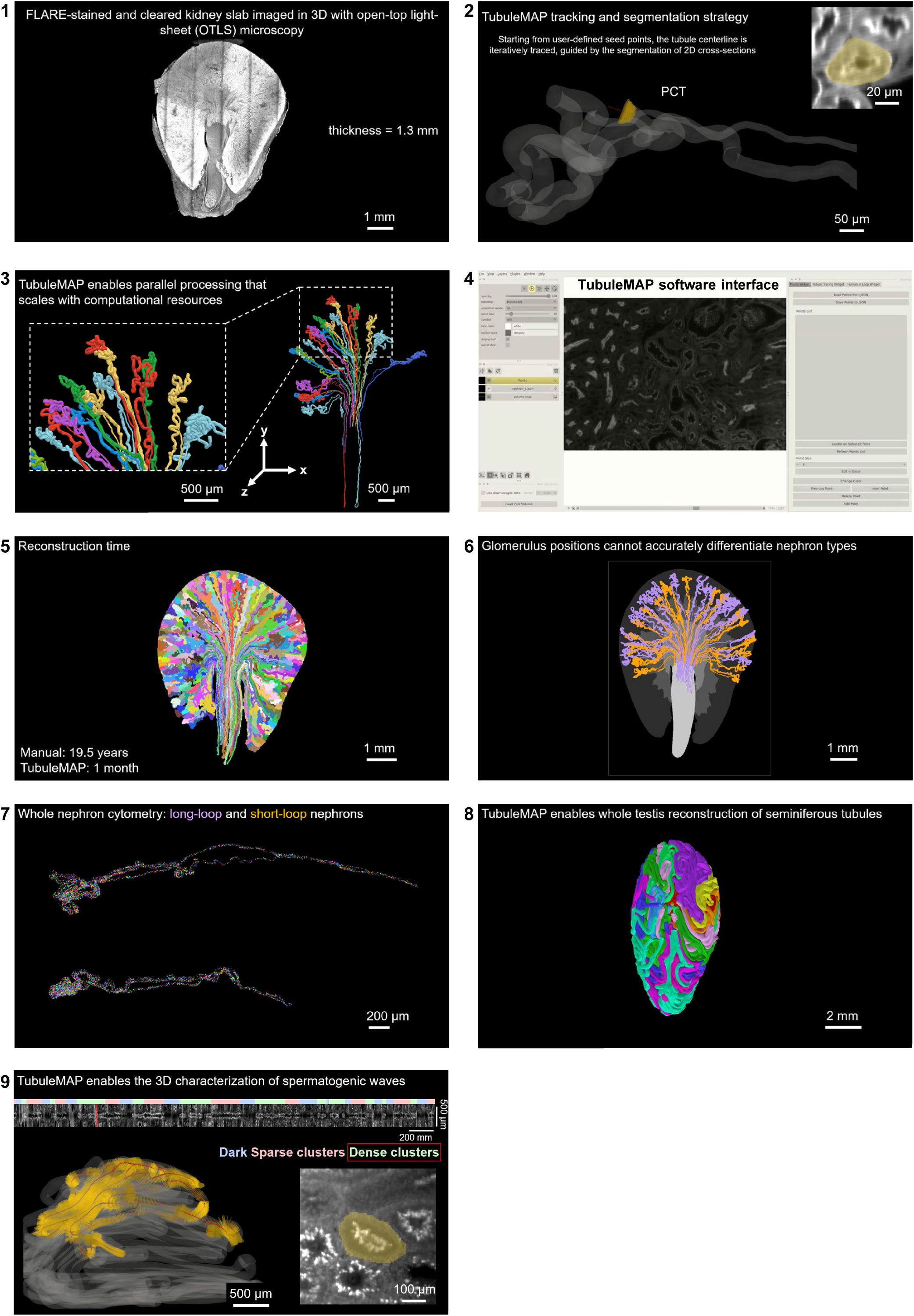
Screenshots of supplementary videos.

### Supplementary Videos

**Supplementary Video 1:** Surface rendering and virtual slices from kidney slab to cropped data with segmented nephron.

**Supplementary Video 2:** TubuleMAP tracking and segmentation strategy.

**Supplementary Video 3:** Parallel processing of tubules trajectories.

**Supplementary Video 4:** TubuleMAP graphical user interface and tracking workflow. Interface is based on napari with multiple widgets for data exploration and human intervention.

**Supplementary Video 5:** Three-dimensional reconstruction of 1000 nephrons from volumetric imaging data.

**Supplementary Video 6:** Spatial distribution of short-loop nephrons and long-loop nephrons across the kidney.

**Supplementary Video 7:** Nuclei distribution in short-loop and long-loop nephrons.

**Supplementary Video 8:** Three-dimensional visualization of seminiferous tubules in cleared testis tissue.

**Supplementary Video 9:** Top: straightened view of the tubule and assignment of spermatogenic wave states. Left bottom: Three-dimensional view of a mouse seminiferous tubule during tracking showing the centerline and orthogonal cross-sections with sampled planes (orange tiles). Right bottom: Raw image views and segmentation masks at the sampled orthogonal plane, along with classified spermatogenic wave states (dark, dense clusters, and sparse clusters).

## Notes

### Competing Interest Statement

Jonathan T C Liu is a cofounder, equity holder, and board member of Alpenglow Bioscience Inc., which has licensed the 3D pathology technologies developed in his lab, including patents related to open-top lightsheet microscopy. Stuart J Shankland is a consultant for Lumen Biosciences. The other authors report no competing interests.

## Reference List

1. Yu ASL, Chertow GM, Luyckx V, Marsden PA, Shkorecki K, Taal MW. Brenner & Rector’s The Kidney. 11th ed: Elsevier; 2020.

2. Kriz W, Kaissling B. Structural Organization of the Mammalian Kidney. Seldin and Giebisch’s The Kidney2008. p. 479–563.

3. Houda A, Shelko N, Bakry MS, Almandouh HB, Micu R, Jankowski PM, et al. Seminiferous tubules and spermatogenesis. Male reproductive anatomy. 2021.

4. Blanc T, Goudin N, Zaidan M, Traore MG, Bienaime F, Turinsky L, et al. Three-dimensional architecture of nephrons in the normal and cystic kidney. Kidney Int. 2021;99(3):632–45.

5. Magri CJ, Fava S. The role of tubular injury in diabetic nephropathy. Eur J Intern Med. 2009;20(6):551–5.

6. Wang Y, Jin M, Cheng CK, Li Q. Tubular injury in diabetic kidney disease: molecular mechanisms and potential therapeutic perspectives. Front Endocrinol (Lausanne). 2023;14:1238927.

7. Lustig L, Guazzone VA, Theas MS, Pleuger C, Jacobo P, Perez CV, et al. Pathomechanisms of Autoimmune Based Testicular Inflammation. Front Immunol. 2020;11:583135.

8. Akhigbe RE, Odetayo AF, Akhigbe TM, Hamed MA, Ashonibare PJ. Pathophysiology and management of testicular ischemia/reperfusion injury: Lessons from animal models. Heliyon. 2024;10(9):e27760.

9. Spangenberg P, Hagemann N, Squire A, Forster N, Krauss SD, Qi Y, et al. Rapid and fully automated blood vasculature analysis in 3D light-sheet image volumes of different organs. Cell Rep Methods. 2023;3(3):100436.

10. Beeuwkes R, 3rd, Bonventre JV. Tubular organization and vascular-tubular relations in the dog kidney. Am J Physiol. 1975;229(3):695–713.

11. Oliver J. Nephrons and Kidneys: A Quantitative Study of Developmental and Evolutionary Mammalian Renal Architectonics.: New York, Hoeber; 1968.

12. Zhai XY, Thomsen JS, Birn H, Kristoffersen IB, Andreasen A, Christensen EI. Three-dimensional reconstruction of the mouse nephron. J Am Soc Nephrol. 2006;17(1):77–88.

13. Christensen EI, Grann B, Kristoffersen IB, Skriver E, Thomsen JS, Andreasen A. Three-dimensional reconstruction of the rat nephron. Am J Physiol Renal Physiol. 2014;306(6):F664–71.

14. Pannabecker TL, Dantzler WH. Three-dimensional architecture of collecting ducts, loops of Henle, and blood vessels in the renal papilla. Am J Physiol Renal Physiol. 2007;293(3):F696–704.

15. Li Y, Cao J, Zhang Q, Li J, Li X, Zhou H, et al. Precise reconstruction of the entire mouse kidney at cellular resolution. Biomed Opt Express. 2024;15(3):1474–85.

16. Torres R, Velazquez H, Chang JJ, Levene MJ, Moeckel G, Desir GV, et al. Three-Dimensional Morphology by Multiphoton Microscopy with Clearing in a Model of Cisplatin-Induced CKD. J Am Soc Nephrol. 2016;27(4):1102–12.

17. Glaser AK, Bishop KW, Barner LA, Susaki EA, Kubota SI, Gao G, et al. A hybrid open-top light-sheet microscope for versatile multi-scale imaging of cleared tissues. Nat Methods. 2022;19(5):613–9.

18. Li Y, Cao J, Li A, Li X, Jiang T, Tuchin VV, et al. Segmentation and reconstruction of renal tubule in mesoscopic mouse kidney images. Seventeenth International Conference on Photonics and Imaging in Biology and Medicine (PIBM 2024)2025.

19. Januszewski M, Kornfeld J, Li PH, Pope A, Blakely T, Lindsey L, et al. High-precision automated reconstruction of neurons with flood-filling networks. Nat Methods. 2018;15(8):605–10.

20. Tavakoli MR, Lyudchik J, Januszewski M, Vistunou V, Agudelo Duenas N, Vorlaufer J, et al. Light-microscopy-based connectomic reconstruction of mammalian brain tissue. Nature. 2025;642(8067):398–410.

21. Stringer C, Wang T, Michaelos M, Pachitariu M. Cellpose: a generalist algorithm for cellular segmentation. Nat Methods. 2021;18(1):100–6.

22. Pachitariu M, Stringer C. Cellpose 2.0: how to train your own model. Nat Methods. 2022;19(12):1634–41.

23. Lee MY, Mao C, Glaser AK, Woodworth MA, Halpern AR, Ali A, et al. Fluorescent labeling of abundant reactive entities (FLARE) for cleared-tissue and super-resolution microscopy. Nat Protoc. 2022;17(3):819–46.

24. Mao C, Lee MY, Jhan JR, Halpern AR, Woodworth MA, Glaser AK, et al. Feature-rich covalent stains for super-resolution and cleared tissue fluorescence microscopy. Sci Adv. 2020;6(22):eaba4542.

25. Klingberg A, Hasenberg A, Ludwig-Portugall I, Medyukhina A, Mann L, Brenzel A, et al. Fully Automated Evaluation of Total Glomerular Number and Capillary Tuft Size in Nephritic Kidneys Using Lightsheet Microscopy. J Am Soc Nephrol. 2017;28(2):452–9.

26. Moore J, Basurto-Lozada D, Besson S, Bogovic J, Bragantini J, Brown EM, et al. OME-Zarr: a cloud-optimized bioimaging file format with international community support. Histochem Cell Biol. 2023;160(3):223–51.

27. Bragantini J, Theodoro I, Zhao X, Huijben T, Hirata-Miyasaki E, VijayKumar S, et al. Ultrack: pushing the limits of cell tracking across biological scales. Nat Methods. 2025;22(11):2423–36.

28. Wang Q, Ding SL, Li Y, Royall J, Feng D, Lesnar P, et al. The Allen Mouse Brain Common Coordinate Framework: A 3D Reference Atlas. Cell. 2020;181(4):936–53 e20.

29. Kriz W, Koepsell H. The structural organization of the mouse kidney. Z Anat Entwicklungsgesch. 1974;144(2):137–63.

30. Jamison RL. Short and long loop nephrons. Kidney Int. 1987;31(2):597–605.

31. Susaki EA, Shimizu C, Kuno A, Tainaka K, Li X, Nishi K, et al. Versatile whole-organ/body staining and imaging based on electrolyte-gel properties of biological tissues. Nat Commun. 2020;11(1):1982.

32. Oakberg EF. A description of spermiogenesis in the mouse and its use in analysis of the cycle of the seminiferous epithelium and germ cell renewal. Am J Anat. 1956;99(3):391–413.

33. Nakata H, Sonomura T, Iseki S. Three-dimensional analysis of seminiferous tubules and spermatogenic waves in mice. Reproduction. 2017;154(5):569–79.

34. Chevalier RL. The proximal tubule is the primary target of injury and progression of kidney disease: role of the glomerulotubular junction. Am J Physiol Renal Physiol. 2016;311(1):F145–61.

35. Nakano T, Nakata H, Kadomoto S, Iwamoto H, Yaegashi H, Iijima M, et al. Three-dimensional morphological analysis of spermatogenesis in aged mouse testes. Sci Rep. 2021;11(1):23007.

36. Nakata H, Wakayama T, Sonomura T, Honma S, Hatta T, Iseki S. Three-dimensional structure of seminiferous tubules in the adult mouse. J Anat. 2015;227(5):686–94.

37. Nakata H, Yamaguchi M, Omotehara T, Ichimura K, Yi SQ, Iseki S. Three-dimensional analysis of seminiferous tubules and spermatogenesis in the musk shrew, Suncus murinus. J Anat. 2025;247(6):1204–14.

38. Walsh CL, Tafforeau P, Wagner WL, Jafree DJ, Bellier A, Werlein C, et al. Imaging intact human organs with local resolution of cellular structures using hierarchical phase-contrast tomography. Nat Methods. 2021;18(12):1532–41.

39. Chen F, Tillberg PW, Boyden ES. Expansion microscopy. Science. 2015;347(6221):543–8.

40. Ku T, Swaney J, Park JY, Albanese A, Murray E, Cho JH, et al. Multiplexed and scalable super-resolution imaging of three-dimensional protein localization in size-adjustable tissues. Nat Biotechnol. 2016;34(9):973–81.

41. Chozinski TJ, Halpern AR, Okawa H, Kim HJ, Tremel GJ, Wong RO, et al. Expansion microscopy with conventional antibodies and fluorescent proteins. Nat Methods. 2016;13(6):485–8.

42. Stringer C, Pachitariu M. Cellpose3: one-click image restoration for improved cellular segmentation. Nat Methods. 2025;22(3):592–9.

43. Pachitariu M, Rariden M, Stringer C. Cellpose-SAM: superhuman generalization for cellular segmentation. 2025.

44. Kim DA, Armenta A, Vaughan JC, Terasaki M, Himmelfarb J, Zheng Y. Hemodynamic simulation and in vitro modeling of three-dimensional glomeruli at anatomical scale. Physics of Fluids. 2025;37(5):051907.

45. Glaser AK, Reder NP, Chen Y, Yin C, Wei L, Kang S, et al. Multi-immersion open-top light-sheet microscope for high-throughput imaging of cleared tissues. Nat Commun. 2019;10(1):2781.

46. Balázs B, Deschamps J, Albert M, Ries J, Hufnagel L. A real-time compression library for microscopy images. bioRxiv. 2017.

47. Hörl D, Rojas Rusak F, Preusser F, Tillberg P, Randel N, Chhetri RK, et al. BigStitcher: reconstructing high-resolution image datasets of cleared and expanded samples. Nature Methods. 2019;16(9):870–4.

